# Cholesterol-mediated activation of the VapC12 toxin modulates growth and drug susceptibility in *Mycobacterium tuberculosis*

**DOI:** 10.1101/2025.10.10.681325

**Authors:** Zohra Hashmi, Shivendra Pratap Singh, Sakshi Talwar, Abhin Kumar Megta, Mitul Srivastava, Debapriyo Sarmadhikari, Sourabh Kundu, Prabhakar Babele, Taruna Sharma, Chandresh Sharma, Shailendra Asthana, Vengadesan Krishnan, Amit Kumar Pandey

## Abstract

Tuberculosis eradication efforts are severely hampered by antimicrobial resistance and bacterial persistence. We previously reported that the VapBC12 toxin–antitoxin (TA) system in *Mycobacterium tuberculosis* (*M. tuberculosis*), regulates disease progression and antibiotic persistence. In this study, we investigated the molecular mechanisms underlying VapC12-dependent modulation of bacterial growth and antibiotic susceptibility. We identified a **C**holesterol **R**ecognition **A**mino acid **C**onsensus (CRAC) motif in the VapC12 toxin that is critical for cholesterol binding and interaction with its cognate antitoxin, VapB12. Heterologous expression of VapC12 in *Mycobacterium smegmatis* induced toxicity, providing a model to investigate its functional activity. We determined the crystal structure of VapB12 at 1.6 Å resolution and used molecular modelling to predict the structure of the VapBC12 complex. Structure–function analyses identified specific residues critical for complex assembly and toxin activity. Mechanistically, our findings suggest that cholesterol activates VapC12 by competitively displacing VapB12 from the CRAC motif, thereby restricting *M. tuberculosis* growth. Finally, we demonstrated that pharmacological inhibition of VapC12 alleviates cholesterol-mediated growth restriction and enhances antibiotic activity. Collectively, these findings establish a mechanistic link between cholesterol sensing, toxin–antitoxin regulation, and antibiotic susceptibility in *M. tuberculosis*, highlighting VapC12 as a potential target for therapeutic strategies to combat tuberculosis persistence.

**Significance Statement:** The rise in MDR and XDR tuberculosis cases poses a critical challenge for disease eradication programs worldwide. This therapeutic failure is driven by the ability of *M. tuberculosis* to generate metabolically dormant “antibiotic persister” cells that are refractory to anti-TB drugs. We previously established that the utilization of host-cholesterol promotes persister formation during infection, a phenotype dependent on the VapC12 ribonuclease toxin. In this study, we decipher the mechanism by which the VapC12 ribonuclease toxin regulates cholesterol-dependent growth restriction and antibiotic tolerance in the host environment. Importantly, we demonstrate that targeting VapC12 as an adjunct therapy significantly enhances the efficacy of anti-tuberculosis drugs by alleviating cholesterol-mediated growth arrest and reducing phenotypic drug tolerance in M. tuberculosis.

## Introduction

A steady rise in the frequency of multidrug-resistant (MDR) and extensively drug-resistant (XDR) tuberculosis continuously challenges global tuberculosis eradication deadlines^1^. The ability of *M. tuberculosis* to persist inside the host, together with the heterogeneity-driven stochastic emergence of a phenotypically tolerant subpopulation, poses a significant challenge in the pursuit of finding a potent anti-TB therapy^2–4^. The diverse microenvironments within granulomas create a high degree of heterogeneity, thereby permitting varying degrees of bacterial growth and fostering the generation of drug-tolerant persisters^3–5^. Consequently, targeting mycobacterial persisters represents a promising strategy to combat antimicrobial resistance and improve tuberculosis treatment outcomes^6–8^.

The generation of the persister in *M. tuberculosis* has been reported to be regulated by intricate signaling pathways, including various transcription factors and toxin-antitoxin (TA) systems, which contribute to metabolic adaptation and bacterial survival under diverse stress conditions such as nutrient deprivation, acidic environment and host immune pressure^9–14^. *M. tuberculosis* possesses a large repertoire of approximately 93 TA modules, classified into five families^15^. Among these, the type II VapBC modules have been the most studied TA family. The VapC toxin typically exhibits RNase activity, modulating growth of *M. tuberculosis* by targeting RNA^16–18^. Although *M. tuberculosis* harbors approximately 50 VapBC modules, only a few have been structurally and biochemically characterized^19–27^, and their roles in mycobacterial pathogenesis remain obscure^13,25^. We have previously demonstrated that host cholesterol utilization is critical for the survival of *M. tuberculosis* during the chronic stage of infection in mice^28^. Subsequently, we showed that VapC12 toxin is essential for the generation of cholesterol-induced persisters in tuberculosis^29^. Interestingly, the strain lacking the *VapC12* gene exhibited a hypergrowth phenotype in both guinea pig and murine infection models, leading to an efficient clearance on antibiotic exposure compared to the wild-type strain^29,30^. This observation aligns with the principle that fast-growing bacteria are more susceptible to anti-TB drugs^31,32^.

To get a mechanistic insight, we developed a surrogate *Mycobacterium. smegmatis (M. smegmatis)* based model system to understand the binding dynamics of the VapBC12 TA system and its implications on the growth regulation of *M. tuberculosis*. We identified a CRAC motif specific to the VapC12 toxin and demonstrated that it overlaps with the antitoxin-binding site. We successfully solved the crystal structure of the VapB12 antitoxin and, by homology modeling, predicted that the VapBC12 complex exists as a hetero-octameric form in a “drone” like structure. We further identified residues from both the toxin and the antitoxin that were critical for the stability and/or activity of the TA system. Substitution of critical residues present in the CRAC motif of the toxin disrupted TA binding, resulting in the generation of constitutively active toxin dimers. Our data suggests that cholesterol activates the VapC12 toxin by competitively displacing the VapB12 anti-toxin from the CRAC motif. Interestingly, we found that binding of both the anti-toxin and cholesterol to the toxin involves distinct residues and that an 11-mer peptide, identified from the interacting interface of TA using computational methods to mimic antitoxin, was sufficient to neutralize the activity of the VapC12 toxin. The peptide-mediated inactivation of the toxin enhanced *M. tuberculosis* growth and potentiated the activity of the anti-TB drugs. The findings of this study may help improve the efficacy of existing anti-tuberculosis regimens by providing a mechanistic basis for developing novel strategies to target drug-tolerant persisters..

## Results

### VapC12 toxin modulates growth in *M. tuberculosis*

We began by validating our earlier findings that cholesterol utilisation modulates the growth of *M. tuberculosis*. For this, we introduced a “replication clock” plasmid, pBP10, that, in the absence of a selectable marker, is lost at a steady, quantifiable rate^33^. The plasmid was electroporated into both the H37Rv and an isogenic strain deficient in the *vapC12* gene (Δ*vapC12)*(Figure 1a). We first assessed the growth of the WT strain in minimal medium supplemented with 0.1% glycerol and 150 µM cholesterol. Bacterial counts were enumerated at different time points for both total and plasmid-containing bacteria. As expected, we observed an increase in the plasmid retention time in cholesterol relative to glycerol media, suggesting a reduced growth rate in cholesterol media (Figure 1b). To further confirm the role of the VapC12 toxin in imparting the above phenotype, we compare growth rates between WT and Δ*vapC12* strains on cholesterol using the above protocol. The absence of the VapC12 toxin in *M. tuberculosis* led to a significant loss of the clock plasmid, resulting in an increased growth rate on cholesterol compared to WT. Consistent with the replication clock data, the bacterial density of the Δ*vapC12* increased 10-fold from day 4 to 8 and 5-fold from day 8 to 12, while that of the WT remained relatively constant (Figure 1c). As a control, we observe no statistically significant difference in bacterial density or plasmid retention time between the WT and Δ*vapC12* strains cultured on glycerol media (Figure S1a). These data provide strong validation of our earlier findings that cholesterol-mediated growth modulation is dependent on VapC12.To further uncover the link between cholesterol and VapC12 toxin, we developed a surrogate model using *M. smegmatis.* Its non-pathogenic nature, shorter generation time, and the absence of the VapBC12 TA locus in its genome made *M. smegmatis* an ideal organism for the current study. Briefly, VapC12 toxin alone or in combination with VapB12 anti-toxin was expressed in *M. smegmatis* strain expressing luminescence protein (Figure 1d). Overexpression of VapC12 toxin restricted the growth of *M. smegmatis* while co-expression of cognate anti-toxin VapB12 restored the growth of the strain to the level of *M. smegmatis* harboring the vector alone in glycerol media, while partial restoration was observed in cholesterol media (Figure 1e). The same was further validated using spot titre on 7H11 agar plates (Figure 1f), and the standard CFU plating experiments (Figure 1Sb-c). A similar phenotype was observed on 7H9 enriched media (Figure S1d-f). Overall, the incomplete rescue by VapB12 in cholesterol-containing medium, compared with full rescue in glycerol-containing medium, supports a carbon-source-dependent difference in VapBC12-mediated growth regulation in the *M. smegmatis* surrogate model (Figure 1e-f and S1b-c). These findings are consistent with cholesterol modulating the functional interaction between VapB12 and VapC12.

**Fig. 1:**
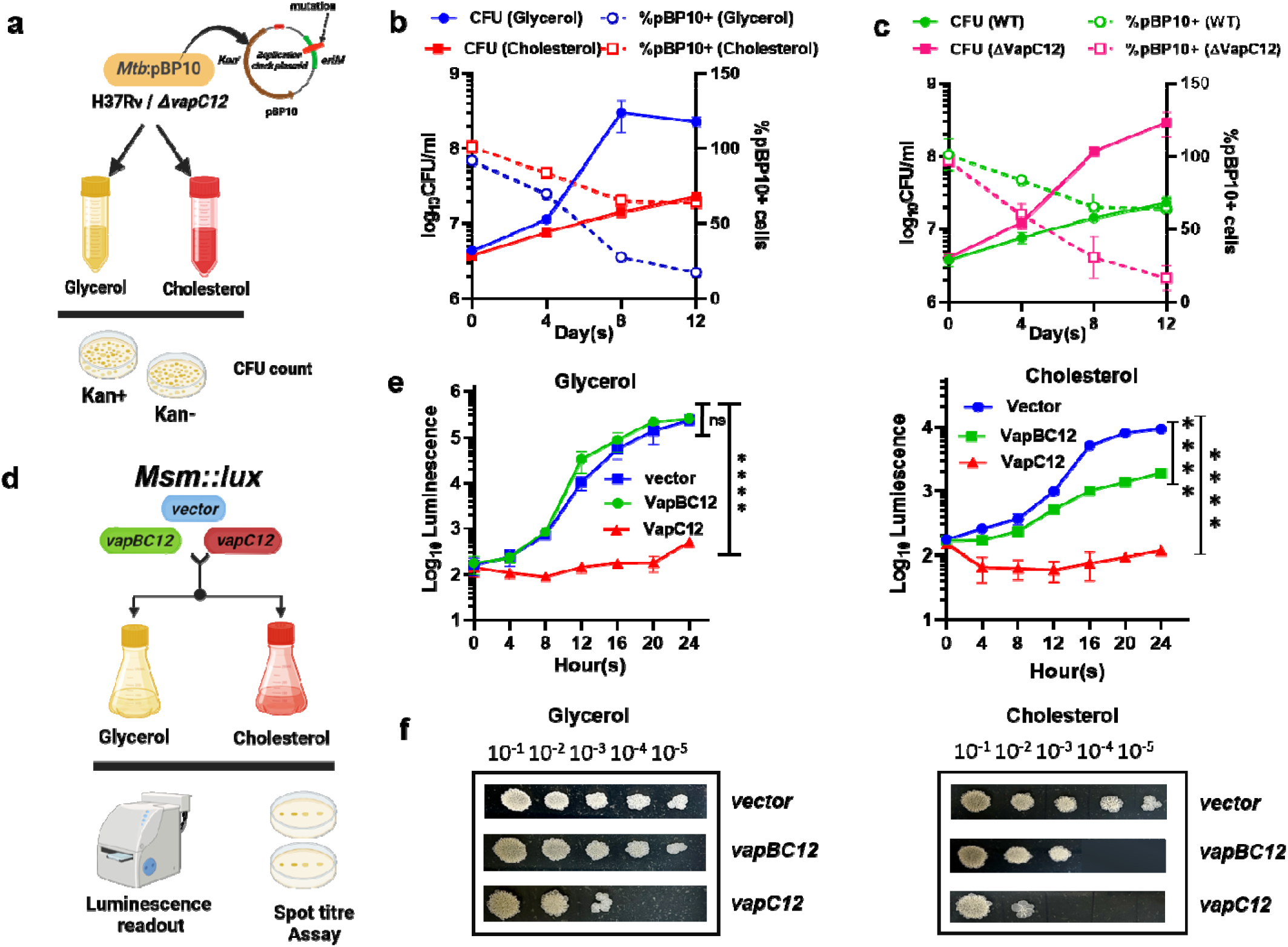
VapC12 toxin-mediated growth modulation in mycobacteria: **(a).** Schematic of the replication clock plasmid (pBP10) assay used to assess replication of wild-type (WT) and ΔvapC12 *M. tuberculosis* in glycerol- or cholesterol-containing media. **(b).** Replication of WT *M. tuberculosis* in glycerol vs cholesterol media. **(c).** Replication of WT *M. tuberculosis* and Δ*vapC12* in cholesterol media measured by plasmid loss over time. (**d).** Schematic showing *M. smegmatis* surrogate strain overexpressing vapBC12 and luciferase. **(e).** Growth analysis of surrogate strains in glycerol and cholesterol media monitored by luminescence. **(f).** Spot titer assay of cultures grown for 24 hours in indicated media. Data represents mean ± SD from three independent experiments. Statistical significance was determined by two-way ANOVA with Tukey’s multiple-comparison test; ns, not significant; ****P < 0.0001.

### Crystal Structure of VapB12 Antitoxin reveals a unique Zn^2+^ binding domain

Our crystallization attempts with VapBC12 from *M. tuberculosis* produced plate-like crystals in a solution containing 0.2 M ammonium chloride-NaOH pH 6.3 and 20% (w/v) PEG 3350 (Figure S2a). These crystals diffracted to a resolution of 1.6 Å and belonged to the tetragonal space group *P*4_1_ (Table 1). Because our attempts to solve the structure by molecular replacement using known toxins and antitoxins as search templates were unsuccessful, we used the intrinsic anomalous signal observed during data processing. Ultimately, the phase calculation by Zn-SAD (zinc single-wavelength anomalous diffraction) yielded interpretable electron density (Table 1). Residues of the N-terminal region (residues 1–44) of VapB12 were clearly visible in the electron density map, whereas its C-terminal region (residues 45–75) and VapC12 were missing. Despite concerted efforts to co-crystallize the VapBC12 complex, only the density corresponding to the N-terminal region of the VapB12 antitoxin was discernible. The crystal structure revealed the DNA-binding domain of VapB12. Four VapB12 molecules (chains A, B, C, and D) in the asymmetric unit form two homodimers (Figure 2a). The two monomers in the dimer interact with each other with a buried interface surface area of ∼1508 Å^2^, mediated by 30 residues via multiple hydrogen bonds and a strong salt bridge between Glu43 and Arg36. The interface between the monomers in the dimer had a complex formation significant score (CSS) value of 1 in the interface analysis by PISA^34^, implying that the interface plays an essential role in complex formation, that is, the functional DNA-binding domain of VapB12. Each monomer displays a typical ribbon-helix-helix (RHH) fold, the common DNA-binding structural motif of antitoxins, with a β1-α1-α2 topology. The β1-strand (residues 2-11) from each monomer forms an antiparallel β-sheet to establish a dimer through multiple interactions, including two strong hydrogen bonds from the main chain of Ala3-Val10 and Ile7-Val5. This β-sheet is flanked by helices α1 (residues 12-24) and α2 (residues 28-43). Helix α2 further stabilizes the dimer through a strong salt bridge between the side chains of Arg36 and Glu43. Moreover, the side chain of Ser30 from helix α2 participates in a hydrogen bond with the main chain of Arg8 from the β1-strand. The basic residue Arg8 likely interacts with DNA, and its presence in the β1-strand classifies the RHH fold in VapB12 as Type II. Similarly, another hydrogen bond interaction between the side chain of Arg21 and the main chain of Pro44 further stabilizes the dimer interface, in addition to nonpolar interactions from multiple residues (e.g., Val5, Ile7, Val10, Leu14, Leu15, Leu18, Phe32, Leu33, Leu34, and Leu37).

**Fig. 2:**
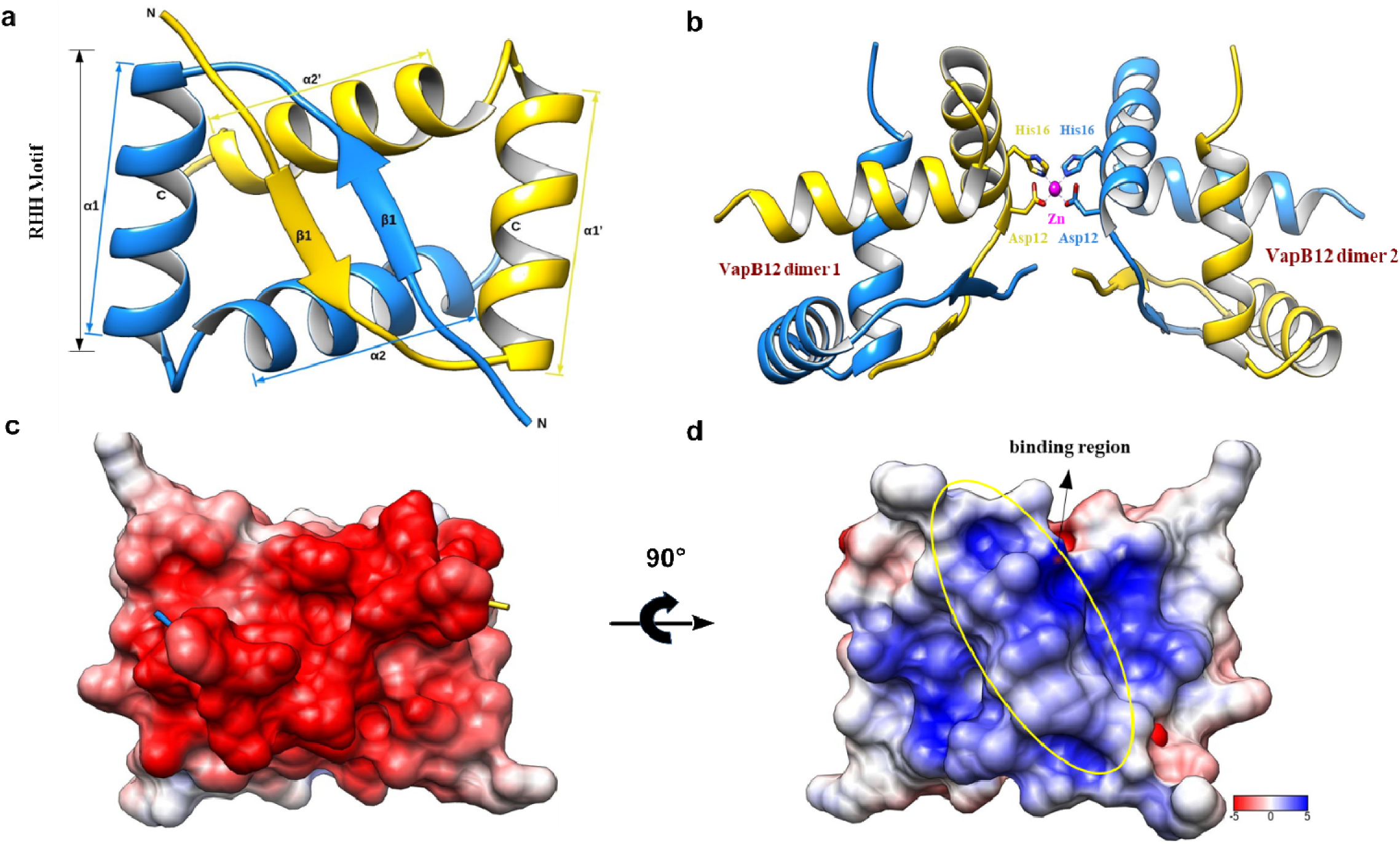
1.6 A° crystal structure of the homodimeric VapB12 antitoxin. **(a).** Monomers of the VapB12 antitoxin are rendered in yellow and blue. The N-terminal DNA-binding ribbon-helix DNA-binding motif (RHH) is labeled. The secondary structural elements (β1, α1, and α2) of VapB12 are labeled on both monomers. **(b).** A cartoon representation of two VapB12 dimers stabilized by Zn coordinated by histidine and aspartic residues. **(c).** The electrostatic surface of the VapB12 homodimer, calculated using APBS and plotted by Chimera. **(d).** View of the VapB12 homodimer with 90° rotation along the x-axis in Fig.2c.

**Table 1.** Data collection and refinement statistics of VapB12 crystallization.

|  | **LrpA<sub>T</sub>** |
| --- | --- |
|  | **Orthorhombic form** |
| **Data collection** |  |
| Space group | *P* 4<sub>1</sub> |
| Cell dimensions |  |
| *a*, *b*, *c* (Å) | 34.66, 34.66, 162.23 |
| *a*, *b*, *g* (°) | 90, 90, 90 |
| X-ray source | ID23-1, ESRF |
| Wavelength (Å) | 0.979507 |
| Resolution range (Å) | 40.55-1.64 (1.66-1.64) |
| No. of unique reflections | 23505 (1174) |
| *R*<sub>merge</sub> | 0.06 (0.79) |
| *R*<sub>*pim*</sub> | 0.02 (0.34) |
| *I*/*s(I)* | 12.6 (2.0) |
| *CC*<sub>1/2</sub> | 0.99 (0.81) |
| Completeness (%) | 99.9 (99.7) |
| Multiplicity | 7.0 (6.3) |
| **Refinement** |  |
| *R*<sub>work</sub> / *R*<sub>free</sub> | 0.172/0.218 |
| Nonhydrogen atoms | 1481 |
| Mean *B* factors ($\text{\AA}^2$) | 38.85 |
| R.m.s. deviations |  |
| Bond lengths ($\text{\AA}$) | 0.011 |
| Bond angles ($^\circ$) | 1.568 |
| Ramachandra plot -favored/allowed/outliers (%) | 100/0/0 |
| PDB ID | 7E4J |

The two dimers interact mainly through a zinc metal ion coordinated by Asp12 and His16 from helix α1 of each dimer, with an average distance of 2.03 Å (Figure 2b). The residue Arg26 from the dimer forms a salt bridge with Glu13 from the adjacent dimer in the crystal lattice. The electrostatic surface of the dimer showed a positive charge on one side and a negative charge on the other side (Figure 2c and 2d). Residues Arg8 (β1-strand), Lys19 (α1-helix), and Arg26 (α2-helix) contribute to the positive charge, while residues Asp31, Glu39, Glu42, Glu43, and Glu46 (α2-helix) and Glu17 (α1-helix) contribute to negative charge. The crystal structures of toxin-antitoxin complexes with DNA (e.g., Arc, MetJ, and CopG) showed an anti-parallel β-sheet from the antitoxin dimer that mediates interactions with DNA by inserting the β-sheet into the major groove of DNA^35–37^. Structural comparison of VapB12 dimers with antitoxin DNA complexes suggests that Arg8 and Qln6 likely mediate interactions with DNA.

### Dimerization of VapC12 is essential for its activity

AlphaFold was used to predict the structure of the TA complex and generate the structure of the VapC12 toxin (Figure 3a-b, S3a,b). Through computational modelling, we identified that VapC12 forms a homodimer via the tryptophan at position 74. To verify this, we expressed His-tagged versions of wild-type and tryptophan-mutated (W_74_G) VapC12 separately in an *E. coli* expression system. Western blot analysis of the recombinant protein revealed that, unlike the wild type, the dimer form of the VapC12 toxin was completely absent from the toxin harbouring the W_74_G point mutation (Figure 3e,S3c,d). Additionally, using the *M. smegmatis* growth model, we demonstrated that toxin oligomerization is essential for its activity (Figure 3f). In sum, our data suggest that the W74 residue is critical for the homo-dimerization of the toxin, which is essential for its ribonuclease activity.

**Fig. 3:**
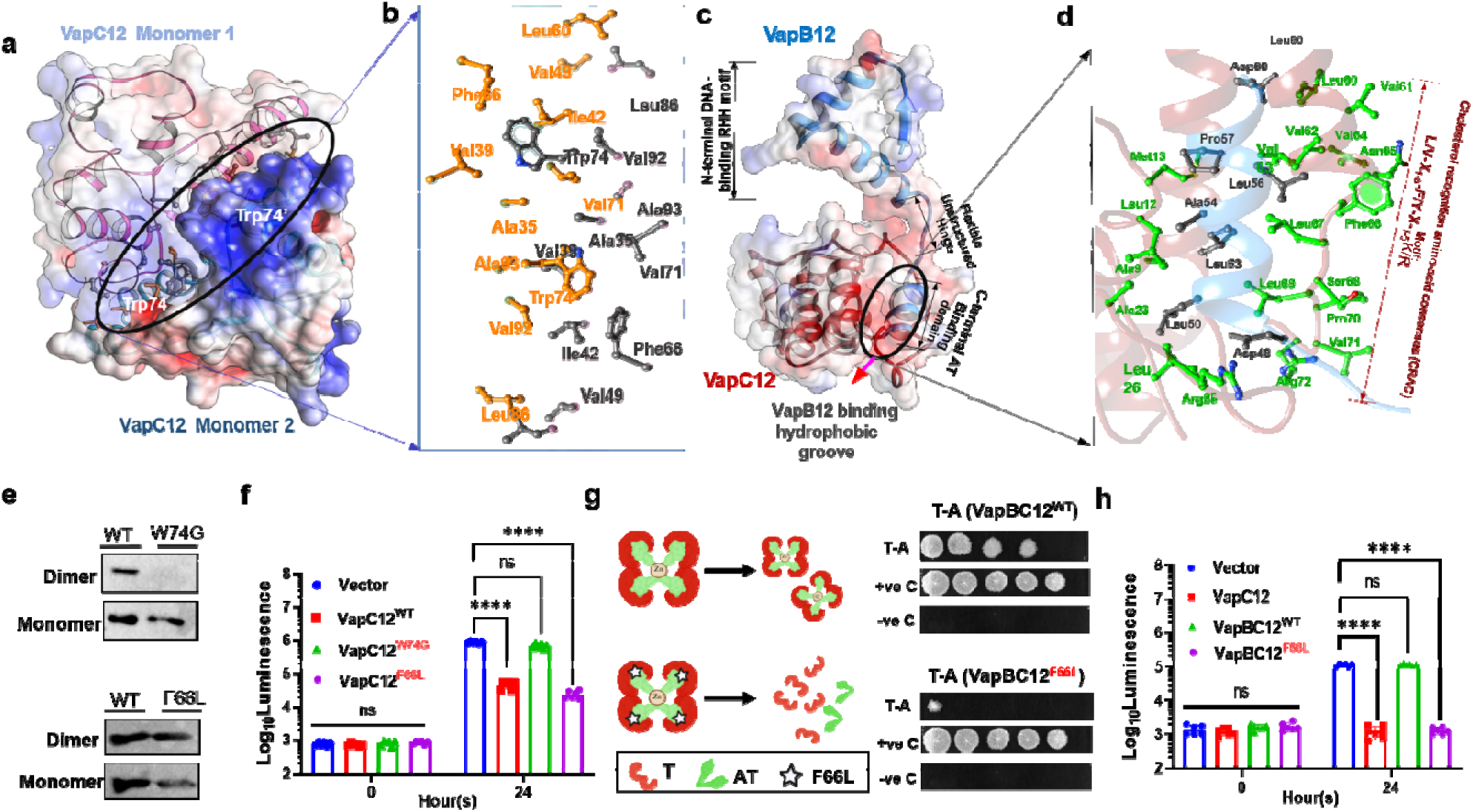
VapC12 is active in homo-dimeric form and interacts with anti-toxin VapB12 via unique CRAC motif: **(a-b):** Structural model of the VapC12 homodimer interface showing hydrophobic interactions centered on Trp74. Interacting residues from the two monomers are shown in gray and orange. **(c-d):** Structural model of the VapBC12 heterodimer interface highlighting ionic and hydrophobic interactions between VapB 2 and VapC12. Residues comprising the CRAC motif in VapC12 are indicated. **(e).** Immunoblot analysis of VapC12 homo-dimerization following glutaraldehyde crosslinking in *E. coli* expressing His-tagged VapC12^WT^, VapC12^W74G^, or VapC12^F66L^. **(f).** Effect of VapC12 mutants on *M. smegmatis* strains expressing vector alone, VapC12^WT^, VapC12^W74G^ and VapC12^F66L^. **(g).** Schematic of the hetero-octameric VapBC12^WT^ and VapBC12^F66L^ complexes and their interaction profiles represented by spot titre in mycobacterial protein fragment complementation (MPFC) assays. **(h)** Viability of *M. smegmatis* strains expressing VapC12, VapBC12^WT^, or VapBC12^F66L^ measured by luminescence. Data in **(f)** and **(h)** represent mean ± SD from three independent experiments. Statistical significance was determined by two-way ANOVA; ns, not significant; **P < 0.01.

### VapC12 interacts with VapB12 through CRAC motif

Sequence analysis of PIN domain-containing prokaryotic toxins identified a cholesterol recognition/interaction amino acid consensus (CRAC) motif uniquely conserved in the mycobacterial VapC12 toxin (Fig. S6). Structural modelling of the VapBC12 complex predicted that the C-terminal region of VapB12 binds a groove in VapC12 overlapping the CRAC motif (Fig. 3c,d), suggesting a direct role for this motif in toxin–antitoxin interaction. To test this, we generated a VapC12 (F66L) mutant targeting a conserved CRAC motif residue. Immunoblot and growth analysis showed that the mutation did not affect toxin homodimerization (Fig. 3e) or VapC12-mediated growth inhibition (Fig. 3f), indicating that the CRAC motif is dispensable for toxin oligomerization and intrinsic activity. However, mycobacterial protein fragment complementation (M-PFC) assays revealed that VapC12 (F66L) failed to interact with VapB12, unlike the wild-type complex (Fig. 3g). Consistently, co-expression of VapB12 was unable to rescue growth inhibition caused by VapC12 (F66L) in *M. smegmatis* (Fig. 3h). Together, these findings demonstrate that residue F66 within the CRAC motif is essential for VapB12 binding and neutralization of VapC12 toxicity.

### Structure-function analysis of the VapBC12 TA system

To define residues critical for VapBC12 complex assembly and function, we generated *M. smegmatis* strains expressing mutant variants of the toxin–antitoxin pair and assessed protein–protein interactions (PPI) using the M-PFC assay. Mutation of Trp74 to glycine in VapC12 (W74G), which disrupts toxin homodimerization, reduced interaction with VapB12 and impaired toxin activity, indicating that toxin oligomerization is essential for both TA complex stability and VapC12 function (Figure 4a, S4a). Similarly, substitution of His16 with alanine in VapB12 (H16A), predicted to disrupt Zn²⁺-mediated antitoxin oligomerization, weakened VapB12–VapC12 interaction in M-PFC assay and compromised rescue of toxin-mediated growth inhibition (Figure 4b, S4a). Consistent with this, M-PFC interaction and growth rescue were observed only in Zn²⁺-supplemented media under the conditions tested, supporting a role for Zn²⁺ in functional VapBC12 association and toxin neutralization (Figure 4c,d, S4b,c). Together, these findings identify Trp74 in VapC12 and His16 in VapB12 as critical determinants of VapBC12 structural integrity and regulation of VapC12 ribonuclease activity.

**Fig. 4:**
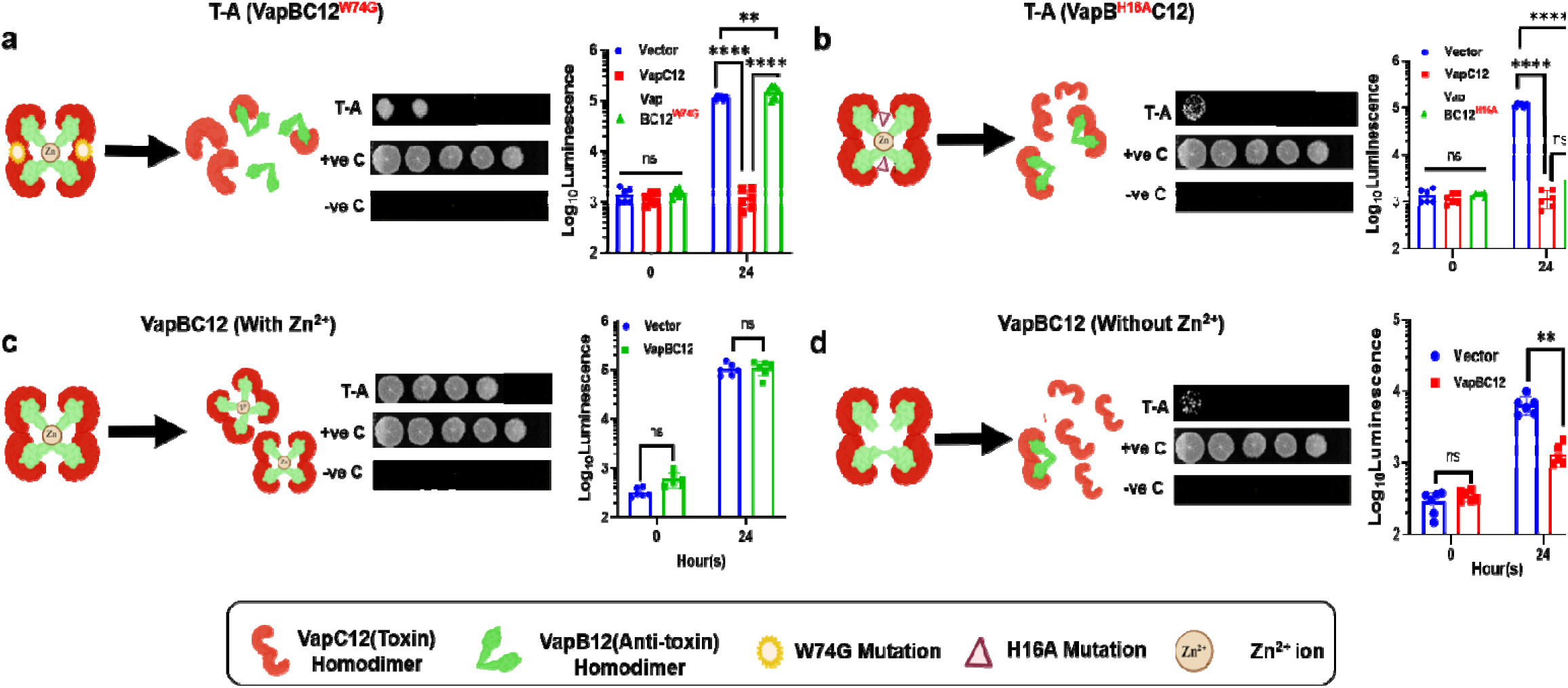
Functional validation of residues critical for TA interaction: **(a)** Schematic and MPFC analysis of the interaction between VapB12 and VapC12^W74G^, followed by growth analysis of *M. smegmatis* strains expressing vector control, VapC12, VapBC12WT, or VapBC12^W74G^. **(b)** Schematic, MPFC analysis, and growth assay of the VapB^H16A12^–VapC12 interaction. **(c,d)** Validation of VapBC12 interaction by MPFC and growth analyses in the presence **(c)** or absence **(d)** of ZnSO4. Agar plates and liquid cultures were prepared in glycerol media with or without zinc supplementation. The interaction between the protein partners was assessed using the MPFC assay by spotting ten-fold serial dilutions of the bacterial cultures onto agar plates supplemented with trimethoprim. Bacterial growth was monitored by constitutive luciferase expression. Luminescence was measured at the indicated time points. Data represent mean ± SD from three independent experiments. Statistical significance was determined by two-way ANOVA with Tukey’s multiple-comparison test; ns, not significant; **P < 0.01; ****P < 0.0001.

### Competitive displacement of VapB12 anti-toxin by cholesterol activates VapC12 toxin

Using an in-silico approach, we demonstrated that cholesterol docks perfectly within the anti-toxin-binding groove formed by the CRAC motif of the toxin (L60-R72) (Figure 5a-b and 3c-d). This suggests that cholesterol may displace the anti-toxin from the CRAC motif of the toxin. We tested this hypothesis with the M-PFC PPI assay. This experiment showed that, unlike glycerol, cholesterol in the medium inhibited binding of the VapB12 anti-toxin to the cognate VapC12 toxin as represented by spot titer(Figure 5c). We expressed C-terminally FLAG-tagged VapC12 alone or the VapBC12 operon in *M. smegmatis* in the presence or absence of cholesterol. Native PAGE followed by anti-FLAG immunoblotting revealed a FLAG-reactive species provisionally assigned to VapBC12 in the absence of cholesterol. Following treatment with 150 μM cholesterol, only a species co-migrating with the VapC12-alone control was detected (Figure 5d). This change is consistent with cholesterol-dependent alteration of VapBC12 association. Interestingly, while co-expression of VapBC12 as an operon partially rescued the VapC12-mediated growth defect in *M. smegmatis* (Figure 1e), inducible expression of the VapB12 antitoxin failed to rescue it in *M. smegmatis* overexpressing VapC12 from a tet-inducible promoter in the presence of cholesterol. In contrast, VapB12 expression effectively restored growth in glycerol- and oleate-containing media (Figure 5e). A concentration-dependent growth inhibition by cholesterol in the strain expressing both toxin and antitoxin further supports the hypothesis (Figure 5f). Additionally, we assessed the binding of 500 nM cholesterol to 50 μg/mL WT-VapC12 and VapC12^F66L^ using BLI. A clear binding association was observed between cholesterol and WT-VapC12, whereas no detectable binding was observed with the VapC12^F66L^ mutant, suggesting that the F66 residue contributes to cholesterol binding (Figure 5g). Overall, the data support our hypothesis that the antitoxin and cholesterol-binding sites overlap at the CRAC motif of the toxin and that dose-dependent competitive displacement of the antitoxin by cholesterol activates the ribonuclease activity of the VapC12 toxin.

**Fig. 5:**
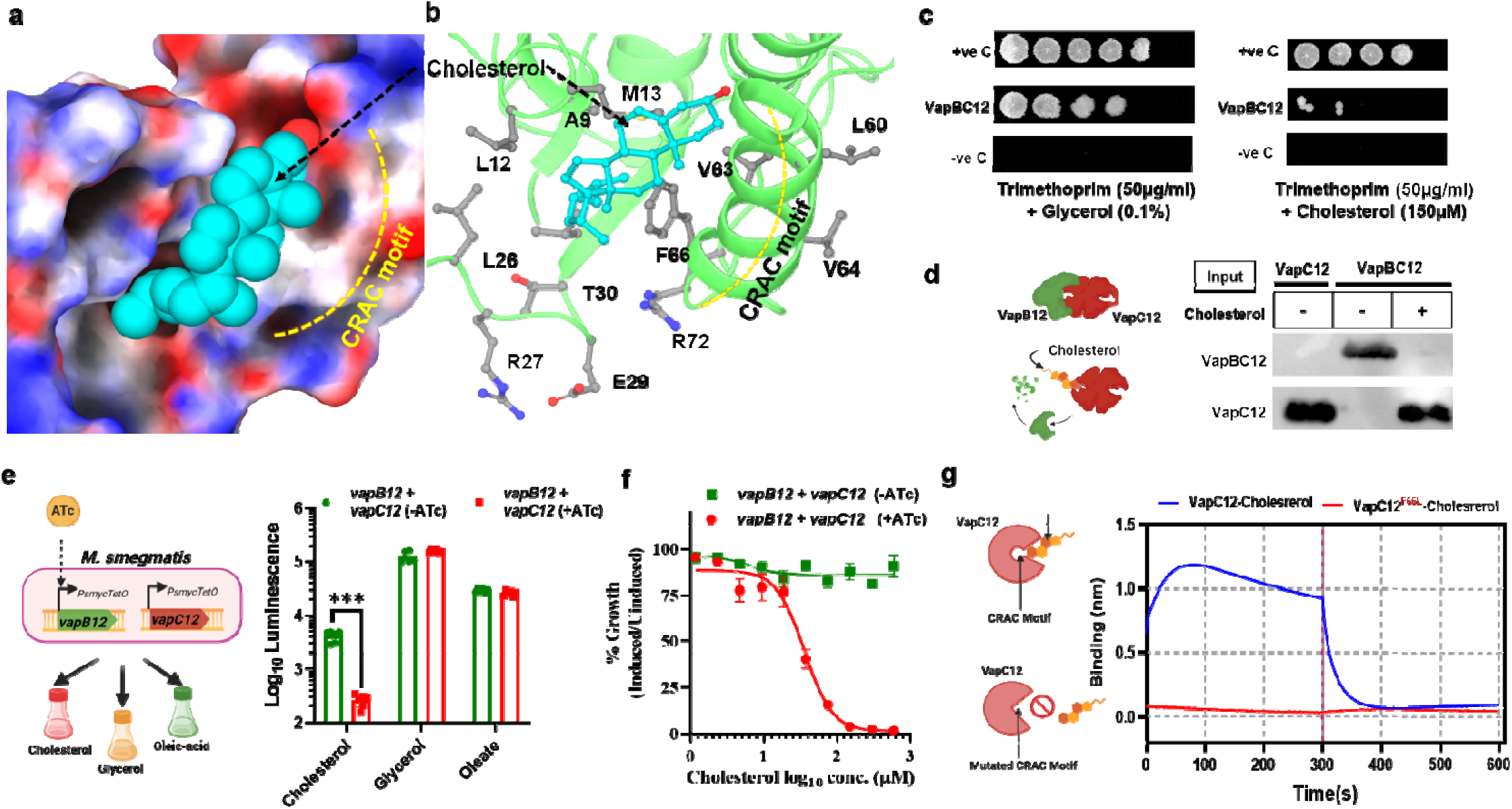
Competitive displacement of VapB12 antitoxin by cholesterol activates VapC12 toxin: **(a,b)** Molecular docking analysis showing the most populated cholesterol-binding pose on VapC12 and residues involved in binding, including the CRAC motif. **(c)** MPFC analysis of VapB12–VapC12 interaction in glycerol- or cholesterol-containing media. Vector alone and ESAT-6/CFP10 served as negative and positive controls, respectively. Growth on trimethoprim-containing 7H11 agar indicates protein–protein interaction. **(d)** Western blot analysis of VapC12 expression in *M. smegmatis* strains expressing C-terminal FLAG-tagged VapC12 alone or VapBC12 as an operon, grown in the presence or absence of cholesterol**. (e)** Growth analysis of *M. smegmatis* strains expressing vapB12 and vapC12 from a tet-inducible promoter in glycerol, cholesterol, or oleate media in the presence of anhydrotetracycline (200 ng/ml). Luminescence was measured after 24 hours of treatment. **(f)** Dose-dependent effect of cholesterol on VapC12-mediated growth modulation in *M. smegmatis*. Bacterial viability was monitored by luminescence. **(g)** BLI sensorgram demonstrating relative binding affinities of 500nM cholesterol to immobilized His-VapC12 or VapC12^F66L^. Data represent mean ± SD from three independent experiments. Statistical significance was determined by two-way ANOVA; ns, not significant; **P < 0.01; ***P < 0.001; ****P < 0.0001.

### Generation and selection of Antitoxin mimicking peptides

Computational analyses identified five VapB12-derived peptides (9–13 aa) from the VapBC12 interface. Protein–peptide docking and MM-GBSA analysis revealed that the 11-aa peptide P2 exhibited the most favorable binding energy (−44.6 kcal/mol) and strong electrostatic complementarity with VapC12 (Figure 6a,b). P2 (L_50_-D_60_) peptide (Figure 6d) docked within the CRAC motif-binding pocket of VapC12, overlapping with the cholesterol-binding site. Docking studies showed that cholesterol and P2 occupied the same binding pocket with comparable binding energies (−43.1 and −44.6 kcal/mol, respectively) (Figure 6c). MD simulations further demonstrated stable binding of both ligands within the pocket (Figure 6e–l). Cholesterol and P2 shared nine interacting residues, including L4, L12, L14, T15, G28, V31, V63, F66, and L67, while both also engaged multiple CRAC motif residues. Cholesterol interacted with V61, V63, N65, F66, and L67, whereas P2 contacted V62, V63, F66, L67, and L69 (Figure 6m). These findings suggest that cholesterol and P2 competitively bind the CRAC motif and may differentially regulate the conformational state of the catalytic PIN domain, thereby modulating VapC12 ribonuclease activity.

**Fig. 6.**
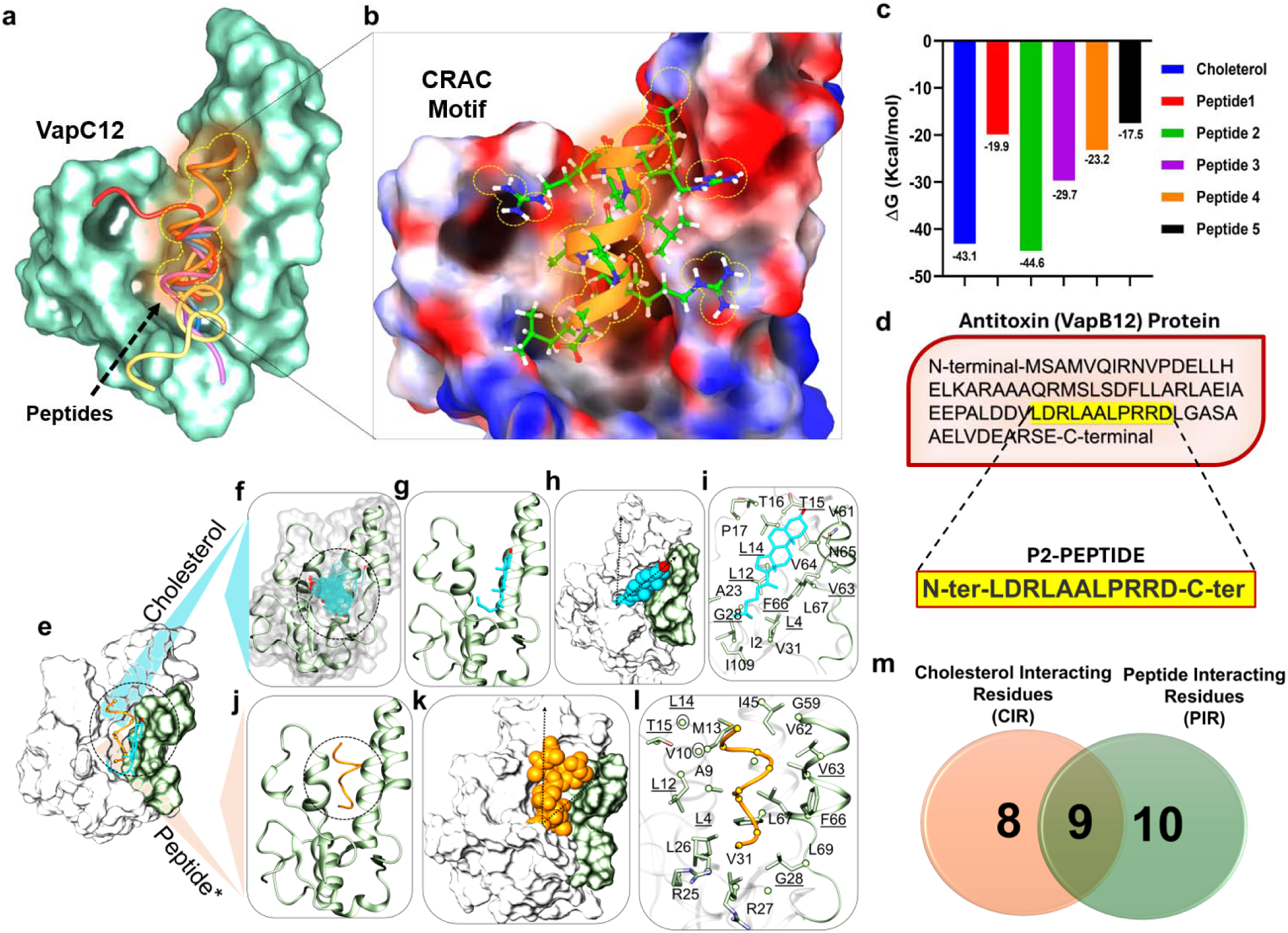
Design and molecular interaction analysis of an antitoxin-mimicking peptide targeting VapC12. **(a)** Predicted binding sites and binding free energies of VapB12-derived peptides (P1–P5) docked onto the VapC12 CRAC motif. **(b)** Structural model of the VapC12–P2 complex showing electrostatic complementarity between P2 arginine residues and the acidic toxin surface. **(c)** Binding free energies of peptides P1–P5 compared with cholesterol. **(d)** Amino acid sequence of peptide P2. (e–i) Molecular interaction analysis of VapC12 with cholesterol and P2. **(e)** Superimposed VapC12–cholesterol–P2 complex with the CRAC motif highlighted. **(f,g)** Docked and best-ranked cholesterol conformations on VapC12. **(h)** Energy-minimized cholesterol-bound structure following molecular dynamics (MD) simulation. **(i)** VapC12 residues within 3.5 Å of cholesterol. **(j,k)** Docked and MD-stabilized conformations of peptide P2 bound to VapC12. **(l)** VapC12 residues within 3.5 Å of peptide P2. Common interacting residues are underlined in **(i)** and **(l).** Binding-axis shifts between cholesterol and P2 are indicated by dotted arrows in **(h)** and **(k). (m)** Venn diagram summarizing shared and distinct VapC12 residues interacting with cholesterol and P2.

### VapC12 toxin inhibition potentiates the activity of the anti-TB drugs

To evaluate inhibition of VapC12 activity, the P2 peptide was synthesized in both L- and D-enantiomeric forms. We first assessed the recovery of intact peptide from *M. tuberculosis* cultures treated with 10 µM of either enantiomer for 4 days (starting at an OD_600_ of 0.01 followed by LC–MS/MS analysis of bacterial lysates. Intact peptide matching the purified standard was recovered exclusively from cultures treated with the D-form, whereas intact L-P2 was not detected in lysates from L-P2-treated cultures under identical conditions, consistent with lower intracellular stability or proteolytic turnover of the L-enantiomer (Fig. S8). Moreover, the L-enantiomer exhibited non-specific effects, promoting bacterial growth in both glycerol- and cholesterol-containing media, possibly due to utilization of the P2 peptide as a nutritional source (Fig. S9a). Consistent with its growth-promoting effect, the L-enantiomer also increased antibiotic susceptibility irrespective of the carbon source, potentially due to enhanced bacterial replication (Fig. S9b–c). In contrast, the L-enantiomer had minimal impact in the THP-1 macrophage model, which may be attributable to its degradation by the proteolytic machinery of the macrophages. (Fig. S9d).

We next investigated whether the D-P2 peptide modulates *M. tuberculosis* growth. Peptide treatment enhanced bacterial growth specifically in cholesterol-containing medium, with no appreciable effect in glycerol-containing medium (Fig. 7a), consistent with a cholesterol-dependent role of VapC12 in growth regulation. Alongside, we also tested the binding of D-P2 peptide with WT- VapC12 and VapC12^F66L^. Similar to Cholesterol, the D-P2 could not bind to VapC12^F66L^ (Fig. 7b). Functional validation using a luminescence-based *M. smegmatis* model showed that the D-form of the P2 peptide restored the growth of VapC12-expressing cells in a concentration-dependent manner in both glycerol- and cholesterol-containing media. Higher peptide concentrations were required in cholesterol-containing media, consistent with competitive binding between cholesterol and P2 at the VapC12 CRAC motif. Importantly, P2 failed to rescue the growth of cells expressing the CRAC mutant VapC12^F66L^, further supporting target-specific activity (Fig. S10a,b).

**Fig. 7.**
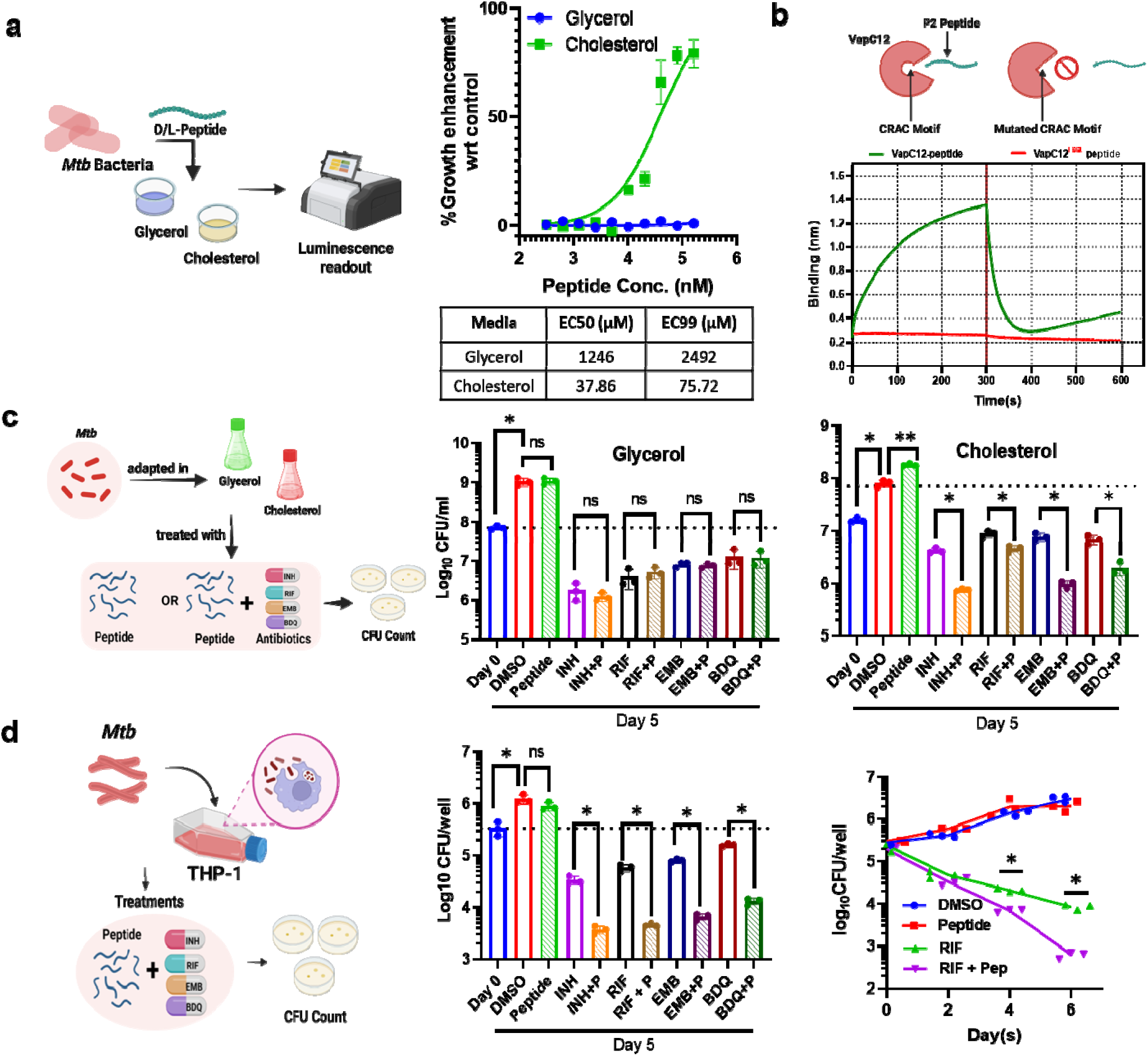
Antitoxin-mimicking peptide P2 potentiates antibiotic killing of *M. tuberculosis*. **(a)** Dose–response analysis of growth enhancement of *M. tuberculosis* in glycerol- and cholesterol-containing media in the presence of D-form P2 peptide normalised with DMSO control. EC₅₀ and EC₉₉ values were calculated by non-linear regression using GraphPad Prism. **(b)** BLI sensorgram demonstrating relative binding affinities of 500nM P2 peptide (D-form) to immobilized His-VapC12 or VapC12^F66L^. **(c)** Survival of *M. tuberculosis* H37Rv treated with 10× MIC isoniazid (INH), rifampicin (RIF), ethambutol (EMB), or bedaquiline (BDQ), alone or in combination with 38 µM P2 peptide in glycerol and cholesterol media. **(d)** Intracellular survival of *M. tuberculosis* in THP-1-derived macrophages infected at MOI 1:1 and treated with 1× MIC antibiotics with or without P2 peptide. Followed by kill kinetics in presence of 1X MIC of RIF alone or in combination of 38 µM P2 peptide. DMSO served as treatment control. Bacterial survival was determined by CFU enumeration. Data represented mean ± SD from three independent experiments. Statistical significance was determined by two-way ANOVA; ns, not significant; **P < 0.01; ***P < 0.001; ****P < 0.0001.

We further examined whether the peptide-mediated growth modulation affected antibiotic susceptibility. Treatment with the VapB12-mimicking P2 peptide in combination with 10× MIC of anti-tubercular drugs resulted in enhanced antibiotic killing and drug potentiation in cholesterol-containing medium, but not in glycerol-containing medium (Fig. 7c), supporting a cholesterol-dependent role of VapC12 in drug tolerance. This phenotype was further recapitulated in the THP-1 infection model. Notably, co-treatment with rifampicin and D-P2 resulted in a more sustained, time-dependent reduction in bacterial viability compared with rifampicin alone, consistent with reduced development or maintenance of drug-tolerant populations (Fig. 7d).

Based on these findings, we designed and patented related peptide molecules (Indian Patent No. 202611034937), which are currently being evaluated by our laboratory in in vitro and animal models of tuberculosis infection. Collectively, these findings demonstrate that the D-form of the P2 peptide targets the VapC12 CRAC motif and modulates cholesterol-dependent VapC12 activity, thereby reducing drug tolerance and enhancing the activity of anti-tubercular therapy against *M. tuberculosis*.

## Discussion

The increased prevalence of immunocompromised individuals, coupled with an increase in the frequency of drug-resistant tuberculosis cases, makes treating this disease even more challenging^39,40^. A treatment regimen for latent TB infection exists^39,40^. However, there is no clinically available drug to inhibit the formation of persister population in *M. tuberculosis*. Recent studies have shown that inhibition of the trehalose catalytic shift^4^, blocking stringent response by hyperphosphorylated guanosine^41^, repression of dihydrolipoamide acyltransferase (DlaT)^42^, an enzyme required to resist nitric oxide-derived reactive nitrogen intermediates, or targeting the cytoplasmic membrane^43,44^ reduces the persister population, resulting in enhanced effectiveness of first-line and second-line anti-TB drugs in macrophage infection models. While these compounds directly kill the persister population, inhibitory molecules targeting the modulatory pathways involved in persister formation have not been explored. Host innate factors, such as cholesterol and other lipids, are important determinants of the mycobacterial phenotype within infected cells^5,28^. Targeting triacylglycerol synthesis in the host mitigates *M. tuberculosis* persistence and thus improves antibiotic clearance in the presence of antibiotics^5^. Several studies have also suggested that targeting the cholesterol metabolism pathway of *M. tuberculosis* may offer a promising therapeutic approach against tuberculosis^45–47^.

Previous studies from our group have demonstrated that cholesterol-mediated modulation of mycobacterial growth and pathogenicity within the host depends on the VapC12 toxin ^28–30^. VapC ribonuclease toxins from type II TA systems in *M. tuberculosis* are usually activated upon the dissociation and subsequent degradation of their cognate antitoxins. Antitoxins usually possess intrinsically flexible domains susceptible to proteolysis, which are mitigated by strong binding to the toxin. While some proteases responsible for anti-toxin degradation have been identified^22,48–50^, the specific dissociation signals that trigger selective proteolysis remain poorly understood. In this study, we employed a multifaceted approach to elucidate the mechanism underlying cholesterol-mediated dissociation of VapB12 and the subsequent activation of the VapC12 toxin in *M. tuberculosis*. Furthermore, as a proof-of-concept, we successfully demonstrated that the VapC12 toxin can be inactivated by a synthetic peptide designed from the VapB12 groove, based on interaction mapping and protein-protein interaction analysis. The most potent peptide (P2) in D form binds the CRAC motif of VapC12 and potentiates the activity of anti-TB drugs in macrophage infection model.

CRAC motifs, which enable cholesterol binding, are common in eukaryotic membrane proteins (e.g., GPCRs) and viral entry proteins^53,54^. These motifs are also found in some bacterial toxins, such as alpha-hemolysin, where they are essential for cholesterol binding and pathogen-associated toxicity^55^. In the current study, we report the presence of a CRAC motif overlapping the antitoxin-binding site in the VapC12 toxin. Our data further suggests that, while binding the antitoxin to the CRAC motif neutralizes its effect, displacement of the antitoxin by cholesterol activates VapC12 ribonuclease activity. However, deeper molecular insights are required to validate this theory.

Our findings corroborate a previous study demonstrating the importance of the hydrophobic core in VapC homo-dimerization^51^. We identified Trp74 as a critical determinant of VapC12 homodimerization and growth-inhibitory activity (Figure 3e-f). Consistent with the structural analysis of type II TA systems in various pathogens^23,24,52^, we predict that the VapBC12 TA system forms a hetero-octamer, which may contribute to toxin neutralization. While divalent metal ions (Mg^2+^, Mn^2+,^ and Ca^2+^) are known to be essential for the catalytic activity of toxin ribonucleases in several VapBC TA systems ^19,24,51^, their role in the stability of the TA system remains unclear. The zinc-mediated contact between VapB12 dimers in the crystal structure, together with the H16A and zinc-supplementation phenotypes, supports a model in which Zn²⁺-mediated antitoxin oligomerization contributes to functional complex assembly and toxin neutralization. The oligomeric states of full-length VapB12 and VapBC12 remain to be established in solution. The metal-binding residues (Asp12 and His16) appear unique to VapB12, as they are not conserved in structural homologs identified by the Dali search (Figure S2B). Whether VapB12 oligomerization influences binding to the TA promoter and transcriptional regulation remains to be explored.

Contrary to the paradigm of TB research that focuses on the development of antimicrobial peptides that activate toxins, to the best of our knowledge, we are the first to demonstrate that a peptide that inactivates VapC12 toxin activity restores bacterial replication under cholesterol conditions and reduces phenotypic drug tolerance in M. tuberculosis, thereby enhancing the effectiveness of anti-TB drugs (Figure 8). Thus, such peptides or small molecules that mimic these peptides hold promise for evaluation as an adjunct to extant TB drugs for the treatment of tuberculosis. Nonetheless, the peptide concentration is too high to be used in an *in-vivo* infection model. We are now focused on peptide modulations and the screening of small compounds for *in-vivo* testing.

**Fig. 8.**
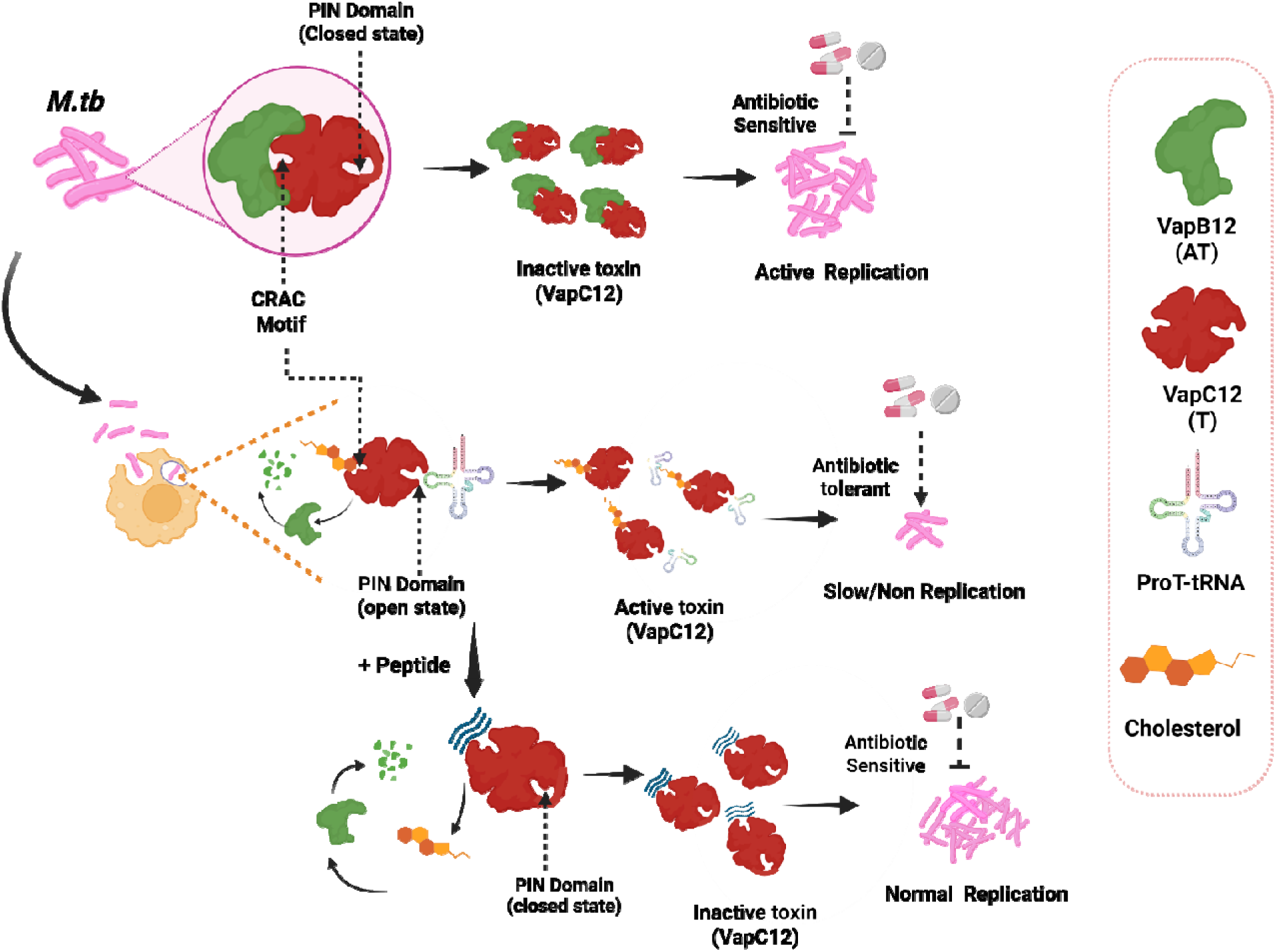
Model for cholesterol-dependent regulation of VapC12 activity in *M. tuberculosis*. Under basal growth conditions, the VapB12 antitoxin binds the CRAC motif of VapC12, maintaining the toxin in an inactive state and permitting bacterial replication. During infection, host-derived cholesterol competitively displaces VapB12 from the CRAC motif, activating VapC12 ribonuclease activity. Activated VapC12 slows bacterial replication and promotes phenotypic antibiotic tolerance. The synthetic antitoxin-mimicking peptide D-P2 competitively binds the CRAC motif, preventing cholesterol-mediated toxin activation, relieving growth restriction, and restoring antibiotic susceptibility.

In conclusion, the presented strategy addresses a major barrier to efficient TB treatment regimens. The role of the VapBC12 TA system in cholesterol-dependent growth modulation and antibiotic tolerance underscores the importance of targeting this TA system during M. tuberculosis infection. Collectively, we identified a mechanism by which cholesterol modulates VapC12 activity to restrict growth and promote drug tolerance, and, as a proof of concept, showed that targeting this pathway as an adjunct therapy potentiates existing anti-tubercular drugs.

## Materials and Methods

### Bacterial strains and growth conditions

*M. smegmatis* mc^2^155 and *M. tuberculosis* strains were cultured on Middlebrook 7H9 broth (Difco, BBL]. Bacterial enumeration was done on 7H11 (Difco, BBL) agar plates supplemented with OADS. *E. coli* XL1 blue and BL21 (DE3) strain was grown in Luria Bertani Agar, Miller [catalog no. M1151-500G] and Luria Bertani Broth [catalog no. M1245-500G]. Strains and primers used in this study are listed in Tables S2, S3, and S4, respectively. Detailed information of bacterial strains and growth conditions can be found in SI Appendix.

### Replication Kinetics studies

The wild-type and VapC12-deleted *M. tuberculosis* strains were transformed with a “molecular clock” plasmid, pBP10 [gift of D. Sherman^33^, which is released from replicating bacteria in the non-selective media. The optimal dilutions of the cultures were plated on 7H11 agar plates, supplemented with or without kanamycin (30 mg/ml), to assess replication rates and growth, respectively. The following formula was used to determine the plasmid retention represented by pBP10+ cells of the two strains in different media:

% pBP10+ cells = CFU on Kan-supplemented plate/CFU on plain plate *100

Detailed protocol is provided in the SI Appendix.

### Growth curve studies

For experiments related to *M. smegmatis* models, Log-phase cultures of wild-type *M. smegmatis* mc^2^155, VapC12, VapB12, and VapBC12 overexpressing strains were washed with 1X PBST and inoculated in 7H9 media supplemented with 0.01% tween 80 (Sigma) and minimal media supplemented with 0.1% glycerol, 150µM cholesterol, and 200 µM oleic acid, at an absorbance of 0.001. Aliquots of the cultures were obtained at different time points, and luminescence was recorded in a 96-well white plate using SpectraMax iD3 Multi-Mode Microplate Readers, and optimal dilutions were prepared to be plated on 7H11 agar plates for bacterial enumeration.

### VapBC12 expression and protein purification

VapB12 (Rv1721c),VapC12 (Rv1720c) and VapC12^F66L^ were expressed and purified as described earlier^24^. Briefly, the two genes were cloned separately into pET28a and transformed into BL21 (DE3). VapC12^F66L^ was generated using SDM generation protocol. The transformed cells were grown overnight, and the culture was inoculated in 1 liter of new LB medium (1:100) supplemented with 100 mg/ml of kanamycin and allowed to grow until the OD_600_ reached 0.6. The culture was induced with 1 mM IPTG and let to grow at 37°C with 120 rpm shaking overnight. Cultured bacteria were harvested by centrifugation at 6,000 × g for 10 min and analyzed for expression by SDS-PAGE. VapB12 was isolated from soluble fraction while VapC12 and VapC12^F66L^ proteins was extracted from inclusion bodies (IBs) in the pellet via urea method. For a detailed protocol, refer to the SI Appendix.

### SDM (Site Directed Mutagenesis) generation

The SDMs in the *vapC12* gene, wherein phenylalanine (F66) and tryptophan (W74) were mutated to leucine (L66) and glycine (G74), respectively, and the vapB12 gene where histidine (H16) was mutated to alanine (A16), were generated by DpnI enzyme (NEB) treatment. Mutants were generated in constructs containing either VapB12 or VapC12, or both VapBC12 together, cloned in a single plasmid as an operon. The cloned constructs were used as templates for PCR amplification with the SDM primers listed in Table S4. Extended methodological information in the SI Appendix.

### Chemical cross-linking

Protein lysates were collected from wild-type and mutated strains of His6-VapC12 in E. coli (DE3) and given a treatment of 0.001% of chemical cross-linker, glutaraldehyde, for 5 min before preparing the sample in Laemmli buffer by boiling at 95° Celsius, separated on a 12% SDS-polyacrylamide gel, and analyzed with western blot using His-tagged antibody. Detailed protocol in the SI Appendix.

### Mycobacterial Protein Fragment Complementation (M-PFC)

The plasmids pMD101 and pMD102 (Kindly provided by Dr. Ashwani Kumar, CSIR-IMTECH, India) containing mDHFR-F(1, 2) and mDHFR-F(3), respectively, were used as vectors to clone VapC12 (toxin) as bait and VapB12 (anti-toxin) as prey protein. SDMs were generated separately in cloned constructs of toxin and anti-toxin using the primers listed in Table S4. F66L and W74G in toxin and H16A in antitoxin. The protocol for *M. smegmatis* culturing was used as described previously^40^.

### Crystallization and structure determination

Crystallization screenings of the VapBC12 complex were performed at 22°C using the hanging-drop vapor-diffusion method, mixing the sample and reservoir solutions in a 1:1 ratio to form a 2 µl drop and equilibrating it against 1000 µl of reservoir solution. A condition containing 0.2 M ammonium chloride-NaOH pH 6.3 and 20% (w/v) PEG 3350 produced diffraction-quality crystals after two weeks (Supplementary Figure 2). The final model revealed only the antitoxin (residues 1-44), and its C-terminal region (residues 45-75) was likely lost during crystallization. After validation, the coordinates of the final model were deposited in the Protein Data Bank (PDB) with accession number (PDB-ID:7E4J). A detailed crystallization protocol can be found in SI Appendix

### Biolayer interferometry (BLI)

The binding of Cholesterol and P2 peptide with his-tagged VapC12 and VapC12^F66L^ were evaluated by BLI using Octet Red96^®^ instrument and Octet® Anti-Penta-HIS (HIS1K) Biosensors (ForteBio, Inc.). Sensors were pre-incubated in MQ water for 10–15 min, and activated with 1X PBS for 600 s. Sensors were incubated with 200 μl of 50 μg/ml VapC12 for 600 s in 1X PBS pH 7.4 to reach levels of 1.5–2.0 nm. Subsequently, the loaded sensors were equilibrated in same buffer (1x PBS) to wash out unbound protein. The VapC12 captured sensors were then dipped in 500nm Cholesterol or P2 peptide (D-form) in the running buffer for the binding assay. The association phase was monitored for 600 s, followed by transfer of the biosensors to ligand-free buffer to monitor dissociation for an additional 600 s. All measurements were performed at 25°C with an agitation speed of 1,000 rpm. Binding kinetics were determined by fitting the association and dissociation sensorgrams using nonlinear regression with a global 1:1 ligand-binding model. Data were analyzed using Octet Analysis Software v10.0 (Sartorius).

### Molecular docking of Cholesterol

The focused docking^64,65^ of cholesterol was performed by taking the centroid of the residues of the CRAC motif. The molecular docking was performed using AUTODOCK4.2 ^66,67^. The most likely pose was selected for energy minimization using molecular dynamics simulation to avoid unfavorable contacts ^68^.

### Free energy analysis through MM-GBSA (molecular mechanics/generalized born surface area)

The average binding energy was calculated by:

ΔGbind=ΔEMM+ΔGsolv + ΔGSA

Where the difference in the minimized energies between ligand and protein complexes is denoted by ΔE_MM_. ΔG_solv_ is the difference in the GBSA solvation energy of the complexes and the sum of solvation energies for the protein and ligand, whereas the differences in surface area energy of the complex and the sum of that in protein and ligand^69^.

### Interaction map-based peptide designing and docking

VapB12 residues that were crucial for binding within the CRAC motif of VapC12 were identified based on interaction types obtained from the VapBC12 (TA) interface, which was then utilized for peptide formation. Based on the interacting interface, the diverse lengths of five peptides were identified from the TA interface. Furthermore, peptide-protein docking was performed using HDOCK (standalone) to obtain the best pose for each peptide. Furthermore, to obtain quantitative values, MM-GBSA calculations were performed for each complex to estimate the binding free energy.

### LC–MS/MS conditions

*M. tuberculosis* cultures (starting OD_600_ 0.01) were treated with 10 µM of either D-P2 or L-P2 peptide for 4 days at 37 °C. Following incubation, bacterial cells were harvested by centrifugation, washed thoroughly with 1× PBS to remove extracellular peptide, and lysed by bead beating in 1X PBS containing 0.01% SDS. Clarified bacterial lysates (50 µg total protein) and purified D- and L-P2 peptide standards (50 µg) were denatured in 8 M urea (1 mg/mL), reduced with 10 mM DTT for 1 h at 37 °C, alkylated with 20 mM iodoacetamide in the dark, diluted to 2 M urea using 50 mM ammonium bicarbonate, and digested with Lys-C (1:20, w/w) overnight at 37 °C to cleave endogenous background proteins while leaving the lysine-free P2 sequence intact. Samples were acidified, desalted, and vacuum-concentrated prior to LC–MS/MS analysis. Approximately 500 ng of each sample was analyzed on a Vanquish™ Neo UHPLC system coupled to an Orbitrap Exploris™ 480 mass spectrometer (Thermo Scientific) equipped with a FAIMS Pro™ interface. Detailed information on LC–MS/MS parameters can be found in the SI Appendix.

### Drug Susceptibility Assessment

The chemically synthesized D- and L-peptide were tested for their effect on *M. tuberculosis* growth, alone or in combination with antibiotics, in glycerol and cholesterol media. The control group received 0.01% DMSO, as the peptide was dissolved in DMSO. *M. tuberculosis* survival was measured by plating the optimal dilutions on 7H11 agar plates supplemented with OADS. Detailed methodology can be found in SI Appendix.

### Macrophage Infection Studies

To assess the effectiveness of AT-peptides on intracellular survival of *M. tuberculosis* in combination with antibiotics, THP-1 monocytes were used. 5×10 ^^5^ cells/well were seeded into a 24-well plate. 50 ng/mL PMA was used for monocyte-to-macrophage maturation. After maturation, the cells were infected with wild-type *M. tuberculosis* at an MOI of 1. After 4 hours post-inoculation, pathogen internalization was measured. The intracellular survival of mycobacteria was measured 4 days post-inoculation. The detailed protocol is provided in the SI Appendix.

### Statistics and reproducibility

The data values are represented as mean ± SD of the replicates from two to three independent experiments. Wherever applicable, the number of replicates per group has been indicated by the dots. GraphPad Prism (ver. 9.0) was used for the statistical analysis of all data, with appropriate tests. Statistical comparisons with *p-values* are mentioned in the respective figure legends.

## Supporting information

This file contains supporting information ias material and methods, figures and tables.

## Author Contributions

Z.H., A.K.P. designed the research; Z.H., S.T., T.S. performed the microbiology-based experiments; C.S. performed the protein purification experiments; S.P.S. and A.K.M. performed crystallography-related experiments; M.S, D.S. and S.A. performed computational studies. Z.H., S.K., and P.B., performed mass spectrometry related experiments. Z.H., S.P.S., S.A., K.V., and A.K.P. analysed the data. Z.H. and A.K.P. drafted the manuscript. All authors contributed to the manuscript revision. A.K.P.: overall supervision.

## Classification

Biological Sciences, Microbiology

## Acknowledgments

The authors acknowledge Translational Health Science and Technology Institute for infrastructural support and the BSL-3 facility staff for assistance with all *M. tuberculosis* experiments. Technical support from Sharad Dwivedi is gratefully acknowledged. The authors thank Vaibhav Kumar Nain for critical editing of figures and legends. The authors also acknowledge the receipt of plasmids and constructs used in this study (Supplementary Information). This work was supported by the Indian Council of Medical Research grant EM/Dev/SG/212/7864/2023 and intramural funding from BRIC-THSTI to A.K.P. Z.H. was supported by a research fellowship from DBTHRDPMU/JRF/BET-21/1/2021-22/236. V.K. acknowledges the Regional Centre for Biotechnology, Faridabad for access to in-house X-ray and central instrumentation facilities, Nicolas Foos for support during data collection, and the DBT-EMBL partnership program for access to the European Synchrotron Radiation Facility. D.S. and S.A. acknowledge support from the NNP grant (NNP-BT/PR40189/BTIS/137/50/2022) and DBT-TRP grant, respectively.

