## Supplementary material for "Cholesterol-mediated activation of the VapC12 toxin modulates growth and drug susceptibility in *Mycobacterium tuberculosis*": This file contains supporting information ias material and methods, figures and tables.

Supporting Information Text

**Materials and Methods**

**Bacterial strains and growth conditions.**

*M. smegmatis* mc^2^155 and *M. tuberculosis* strains were cultured on Middlebrook 7H9 broth (Difco, BBL]. Bacterial enumeration was performed on 7H11 (Difco, BBL) agar plates supplemented with OADS for *M. tuberculosis*, and colonies were counted after 4 days for *M. smegmatis* and 21 days for *M. tuberculosis*. When necessary, Middlebrook medium was supplemented with KAN (25 μg/ml) (Sigma), HYG (50 μg/ml) (Sigma), TRIM (50 μg/ml) (Sigma), and ATc (400 ng/ml) (Cayman), unless otherwise stated. *E. coli* XL1 blue and BL21 (DE3) strain was grown in Luria Bertani Agar, Miller [catalog no. M1151-500G] and Luria Bertani Broth [catalog no. M1245-500G] supplemented with KAN (50 μg/ml) or HYG (150 μg/ml). Strains and primers used in this study are listed in Tables S2, S3, and S4, respectively.

**Replication Kinetics studies**

The wild-type and VapC12-deleted *M. tuberculosis* strains were transformed with a “molecular clock” plasmid, pBP10 [gift of D. Sherman^33^, which is released from replicating bacteria in the non-selective media. The plasmid loss in the absence of the selection marker allows direct measurement of replication, calculated as the ratio of plasmid-carrying bacteria to the total number of bacteria. The VapC12 Comp strain was not used in this assay due to a conflict with the selectable markers. Log-phase cultures of both strains grown in Kan-supplemented Middlebrook 7H9 broth were washed with phosphate-buffered saline containing 0.05% tyloxapol (Sigma) (PBST) to remove residual nutrients. These washed cultures were used to set an OD of 0.01 in minimal media supplemented with 0.1% glycerol (Sigma), referred to as glycerol media, and 150µM cholesterol (Sigma), referred to as cholesterol media. The minimal media was also supplemented with 0.05% tyloxapol (Sigma) to prevent clumping, which could affect the accuracy of the bacterial counts. The optimal dilutions of the cultures were plated on 7H11 agar plates supplemented with and without kanamycin (30 mg/ml) to assess replication rates and growth, respectively. The following formula was used to determine the plasmid retention represented by pBP10+ cells of the two strains in different media:

% pBP10+ cells = CFU on Kan-supplemented plate/CFU on plain plate *100

**SDM (Site Directed Mutagenesis) generation**

The SDMs in the *vapC12* gene, wherein phenylalanine (F66) and tryptophane (W74) were mutated to leucine (L66) and glycine (G74), respectively, and the vapB12 gene where histidine (H16) was mutated to alanine (A16), were generated by DpnI enzyme (NEB) treatment. Mutants were generated in constructs containing either VapB12 or VapC12, or both VapBC12 together, cloned in a single plasmid as an operon. The cloned constructs were used as templates for PCR amplification with the SDM primers listed in Table S4. The PCR product was purified and treated with DpnI enzyme for 4 h at 37°C. No-template and no-DpnI-treatment control samples were also maintained. All the reactions were transformed into *E. coli* XL-1 Blue-competent cells. The constructed products were sequenced and electroporated into *M. smegmatis* -competent cells for further studies.

**Chemical cross-linking**

The vector pET28a was used for the expression of His6-VapC12 in E. coli (DE3) strain. The SDM mutant (W74G) was generated using Dpn1 digestion technique as stated above. The overnight culture was inoculated into 10 milliliters of fresh LB media (1:100) supplemented with 100mg/ml of kanamycin and allowed to grow until the OD_600_ reached between 0.6-0.8. The culture was then induced with 1 mM IPTG (isopropyl-β-D-thiogalacto-pyranoside) (Sigma) and allowed to grow at 37°C for 4 hours with vigorous shaking (120 rpm) before harvesting cells (6 min, 10,000 rpm) and resuspension in 50 mM Tris (pH 8.0), 500 mM NaCl, 5 mM MgCl2, 5 mM β-mercaptoethanol, 10 mM imidazole (Sigma). Cells were disrupted by sonication on ice with 30% amplitude and pulses of 10s : 10s (i.e. energy for 10s followed by a pause for 10s) for about 2 min. Protein lysates were collected from WT and mutated strains and given a treatment of 0.001% of chemical cross-linker, glutaraldehyde for 5 min before preparing the sample in Laemmli buffer by boiling at 95^0^ Celsius, separated on a 12% SDS-polyacrylamide gel and analyzed with western blot using his-tagged antibody.

**Mycobacterial Protein Fragment Complementation (M-PFC)**

*M. smegmatis* cells were independently transformed with M-PFC plasmids producing [F1,2]/[F3] as negative control, Esat-6[F1,2]/Cfp-10[F3] as positive control, and VapC12[F1,2]/VapB12[F3], VapC12:F66L [F1,2]/VapB12[F3], VapC12:W74G [F1,2]/VapB12[F3] VapC12[F1,2]/VapB12:H16A[F3] (Table S2). The transformants were streaked onto a 7H11 medium containing Trimethoprim 50µg/ml along with positive and negative control. The streaked plate was incubated at 37°C. The plates were observed for growth after 4 days.

**VapBC12 expression and protein purification**

VapB12 (Rv1721c) and VapC12 (Rv1720c) were expressed and purified as described earlier^24^. Briefly, the two genes were cloned separately into pET28a and transformed into BL21 (DE3). The transformed cells were grown overnight, and the culture was inoculated in 1 liter of new LB medium (1:100) supplemented with 100 mg/ml of kanamycin and allowed to grow until the OD_600_ reached 0.6. After that, the culture was induced with 1 mM IPTG and let to grow at 37°C with 120 rpm shaking overnight. Cultured bacteria were harvested by spinning at 6,000 *g for 10 min and checked for expression by SDS-PAGE. Majority of the VapC12 (Rv1720c) protein was found as inclusion bodies (IBs) in the pellet. Using sonication and many washing stages, pure IBs carrying rRv1720c were isolated. In brief, one milliliter of pure rRv1720c IBs was dissolved in nine milliliters of buffer (50 mM Tris-HCl [pH 8.0], 300 mM NaCl, 10 mM β-mercaptoethanol, and 8 M urea) and let stand for an hour at room temperature. The sample was then centrifuged at 15,000*g for twenty minutes at 10°C. The obtained supernatant was used to purify recombinant VapC12 protein by immobilized metal ion affinity chromatography using a HisTrap FF column (GE Healthcare Buckinghamshire, UK) under denaturing conditions Protein was eluted using buffer (50 mM Tris-HCl [pH 8.0], 300 mM NaCl, 10 mM β-mercaptoethanol, 8 M urea, and 250 mM imidazole). Denatured purified protein was refolded by dilution in a pulsatile manner at 4°C while being constantly stirred in a refolding buffer (100 mM phosphate buffer [pH 6.4], 300 mM NaCl, and 5 mM β-mercaptoethanol). The refolded target protein sample was centrifuged at 24,000 g for thirty minutes at 4°C. The supernatant including the refolded active protein, was then concentrated and dialyzed three times against the buffer, which consisted of 100 mM phosphate buffer [pH 6.4], 300 mM NaCl, and 5 mM anhydroethanol. The protein's final buffer exchange was carried out using a PD10 desalting column (GE Healthcare Buckinghamshire, UK) using the following parameters: 100 mM phosphate buffer [pH 6.4], 300 mM NaCl, and 10% glycerol. VapB12 (Rv1721c) was purified from the supernatant of lysed-induced cultures using a HisPur Cobalt purification kit (3 ml; Thermo Scientific, catalog no. 90092). Proteins were quantitated by a bicinchoninic acid assay (Thermo Scientific Pierce, Rockford, IL), analyzed and confirmed by SDS-PAGE and Western blotting with anti-His monoclonal antibody (Cell Signaling Technology, Inc., Danvers, MA).

**Crystallization and structure determination**

The toxin (VapC12) and antitoxin (VapB12) were mixed at an equimolar ratio and incubated at 4°C for an hour with gentle shaking. The toxin-antitoxin complex sample was then concentrated to 900 µL, centrifuged at high speed to remove any precipitates, and injected into a HiLoad 16/60 Sephacryl S100 HR (Cytiva) size-exclusion chromatography (SEC) column. The eluted protein fractions were combined, concentrated to 3 mg/ml, and subjected to crystallization trials. Crystallization screenings of the VapBC12 complex were performed at 22°C using the sitting-drop vapor-diffusion technique with an automated liquid handling robotic system (Mosquito, TTP Labtech) against various commercial screening solutions. Protein crystals appeared in a few conditions, with poor diffraction quality. The initial conditions were subsequently reproduced and optimized by hanging-drop vapor-diffusion method using hanging-drop vapor diffusion method by mixing sample and reservoir solution in a 1:1 ratio for making 2 µl drop and equilibrating it against 1000 µl reservoir solution. A condition containing 0.2 M ammonium chloride-NaOH pH 6.3 and 20% (w/v) PEG 3350 produced diffraction-quality crystals after two weeks (Supplementary Figure 2).

Preliminary X-ray diffraction analysis and screening were performed at the home source with a Xenocs GeniX3D Cu (high-flux) microbeam X-ray generator and a MAR 345 image-plate detector (MAR Research). High-resolution diffraction data were collected at the synchrotron beamline ID23-1 (ESRF synchrotron facility at Grenoble, France) (Table 1) using 30% ethylene glycol as a cryoprotectant. Diffraction data were indexed and integrated with *XDS* ^56^ and scaled using *AIMLESS* ^57^ in the *autoPROC* package ^58^. The crystals diffracted to 1.6Å resolution and belonged to the tetragonal space group *P*4_1_, with unit cell parameters of a = 34.66 Å, b = 34.66 Å, c = 162.23 Å, and α = β = γ = 90.00°.

Initial attempts to determine the phases via molecular replacement using the related VapBC structures were unsuccessful. Fortunately, significant anomalous signals were observed during data processing, likely from zinc bound to the protein. We then proceeded with phase determination and preliminary model building via single-wavelength anomalous dispersion (SAD) phase calculation using ShelxCDE ^59^. Subsequent model building with phenix.autobuild ^60^ and the final model refinement cycles through an iterative process utilizing Refmac ^61^ and Coot were performed within the CCP4 ^62^. The refinement process included an initial round of rigid-body refinement, followed by subsequent cycles incorporating TLS parameters determined via the TLSmd server^63^. The final refined model of VapB12 exhibited Rwork/Rfree values of 0.172/0.218, showing excellent geometric properties, with no residues falling within the disallowed regions of the Ramachandran Plot. The final model revealed only the antitoxin (residues 1-44), and its C-terminal region (residues 45-75) was likely lost during crystallization. After validation, the coordinates of the final model were deposited in the Protein Data Bank (PDB) with accession number (PDB-ID:7E4J).

**LC–MS/MS conditions**

Peptides were loaded onto a PepMap™ Neo trap cartridge (100 Å, 5 µm C18, 0.3 × 5 mm) and separated on an Easy-Spray™ PepMap™ Neo analytical column (2 µm C18, 75 µm × 150 mm) at 300 nL/min using 0.1% formic acid in water (A) and 0.1% formic acid in 80% acetonitrile (B), with a gradient of 6% B for 2 min, 6–31% B over 58 min, 31–44% B over 30 min, and 100% B for 9 min. Data-dependent acquisition was performed following calibration with FlexMix (Pierce) using nano-ESI at 2.0 kV, ion transfer tube temperature of 280 °C, FAIMS compensation voltage of −50 V, and RF lens at 40%. Full MS scans were acquired at 60,000 resolution over m/z 350–1200, and MS/MS spectra were acquired at 15,000 resolution using HCD at 30% collision energy for precursor ions (>5.0 × 10³ intensity; charge states 2+–6+), with a dynamic exclusion of 45 s. Protein identification was performed using Proteome Discoverer with the Sequest algorithm against a manually curated database, allowing up to two missed cleavages, a precursor mass tolerance of 10 ppm, carbamidomethylation of cysteine as a static modification, methionine oxidation as a dynamic modification, and a target false discovery rate of 1%.

**Drug Susceptibility Assessment**

The chemically synthesised D-peptide of Anti-toxin was tested on *M.tb* growth, alone or in combination with antibiotics, in glycerol and cholesterol media. The log-phase culture of wild-type *M.tb* grown in 7H9 media supplemented with OADS and 0.01% Tween 80 was washed with PBST to remove remaining nutrients. This was diluted to 0.01 OD in glycerol and cholesterol media to adjust the growth environment. Once the culture reached an OD of 0.2, it was given the treatment of peptide alone or in combination with antibiotics. The control group was given treatment with 0.01% DMSO as the peptide was dissolved in DMSO. After 4 days of treatment, *M.tb* survival was measured by plating the optimal dilutions on 7H11 agar plates supplemented with OADS. For time-dependent kill kinetics, CFU was measured at indicated time points.

**Macrophage Infection Studies**

To assess the effectiveness of AT-peptides on *M.tb* intracellular survival in combination with antibiotics, THP-1 monocytes were used. 5*10^5^ cells/well were seeded on a 24-well plate. 50 ng/ml PMA was used for monocytes to macrophages maturation. After maturation, the cells were infected with wild-type *M.tb* at an MOI of 1. Past 4 hours, infection establishment was measured. Briefly, cells in three replicative wells were washed thrice with 1X PBS (molecular grade) to remove extracellular bacteria. Later, it was treated with PBS supplemented with 0.01% Triton-X100 to lyse the cells. The other wells were given treatment with peptide alone or in combination with antibiotics. The control group was maintained as mentioned above. After 4 days of treatment, the intracellular survival of mycobacteria was measured by lysing the cells followed by plating of cell lysates at optimal dilutions onto 7H11 agar plates supplemented with OADS.

Figures

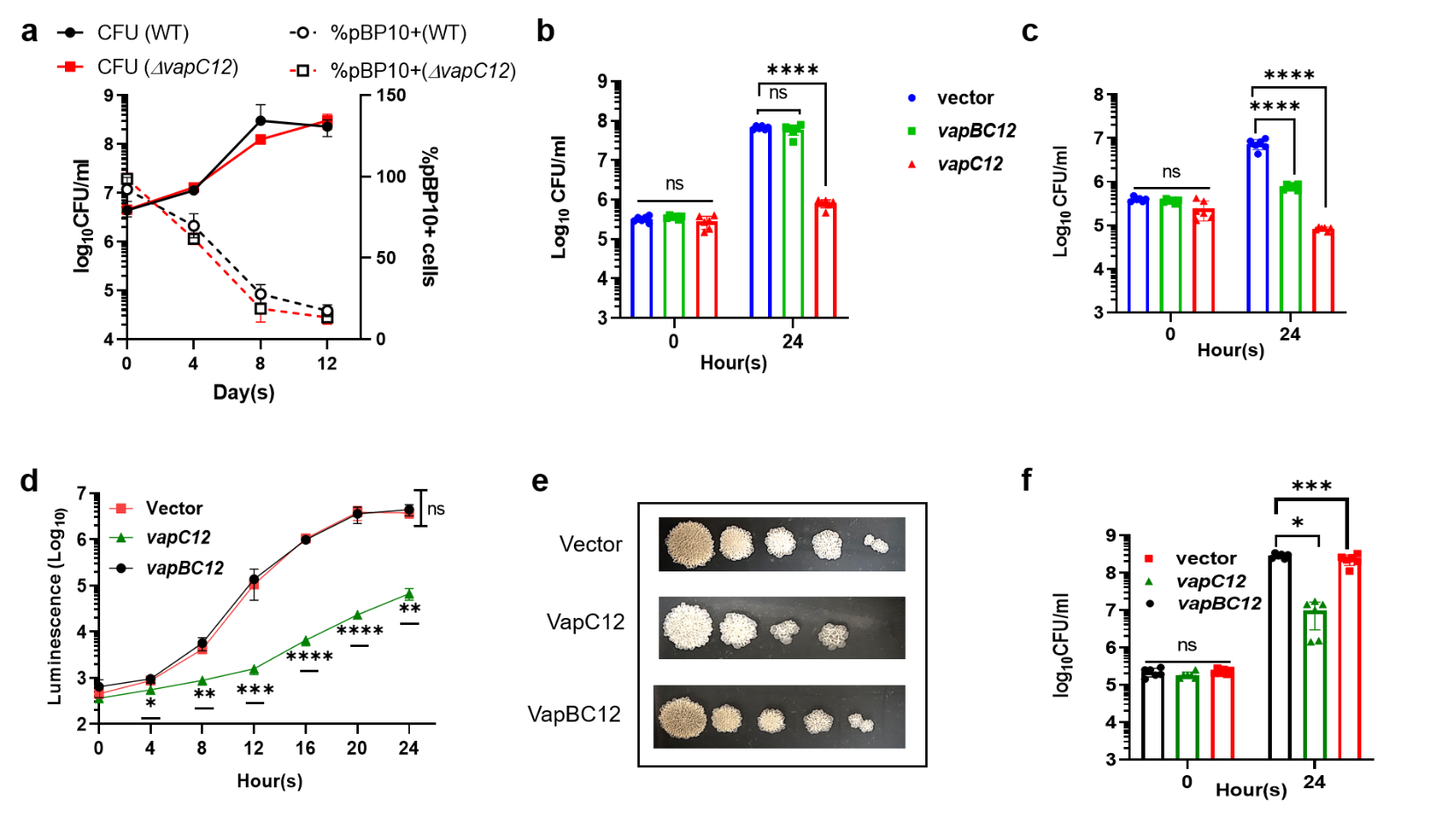

**Fig.S1: VapC12-mediated growth modulation in mycobacteria.** (a). Dual-axis graph of bacterial growth (left y-axis, represented by solid lines) and % pBP10+ cells, plasmid retention (right y-axis, represented by dotted lines) of wild-type and ∆*vapC12* *M. tuberculosis* in glycerol media. (b-c). *In-vitro* growth analysis of *M. smegmatis* expressing vector only control, VapC12, and VapBC12 using CFU enumeration in glycerol (b) and cholesterol media (c). (d-f). *In-vitro* growth analysis of the above-mentioned *M. smegmatis* strains in 7H9 media, represented by luminescence (d), spot titer assay (e), and CFU enumeration (f). The experiment was performed in triplicate with two technical replicates within each biological replicate, and the data plotted represent mean ± the SD. Comparisons for statistical significance have been made using two-way ANOVA test. ns, non-significance,*,p<0.05 **, p < 0.01, ***, p <0.001, ****, P<0.0001.

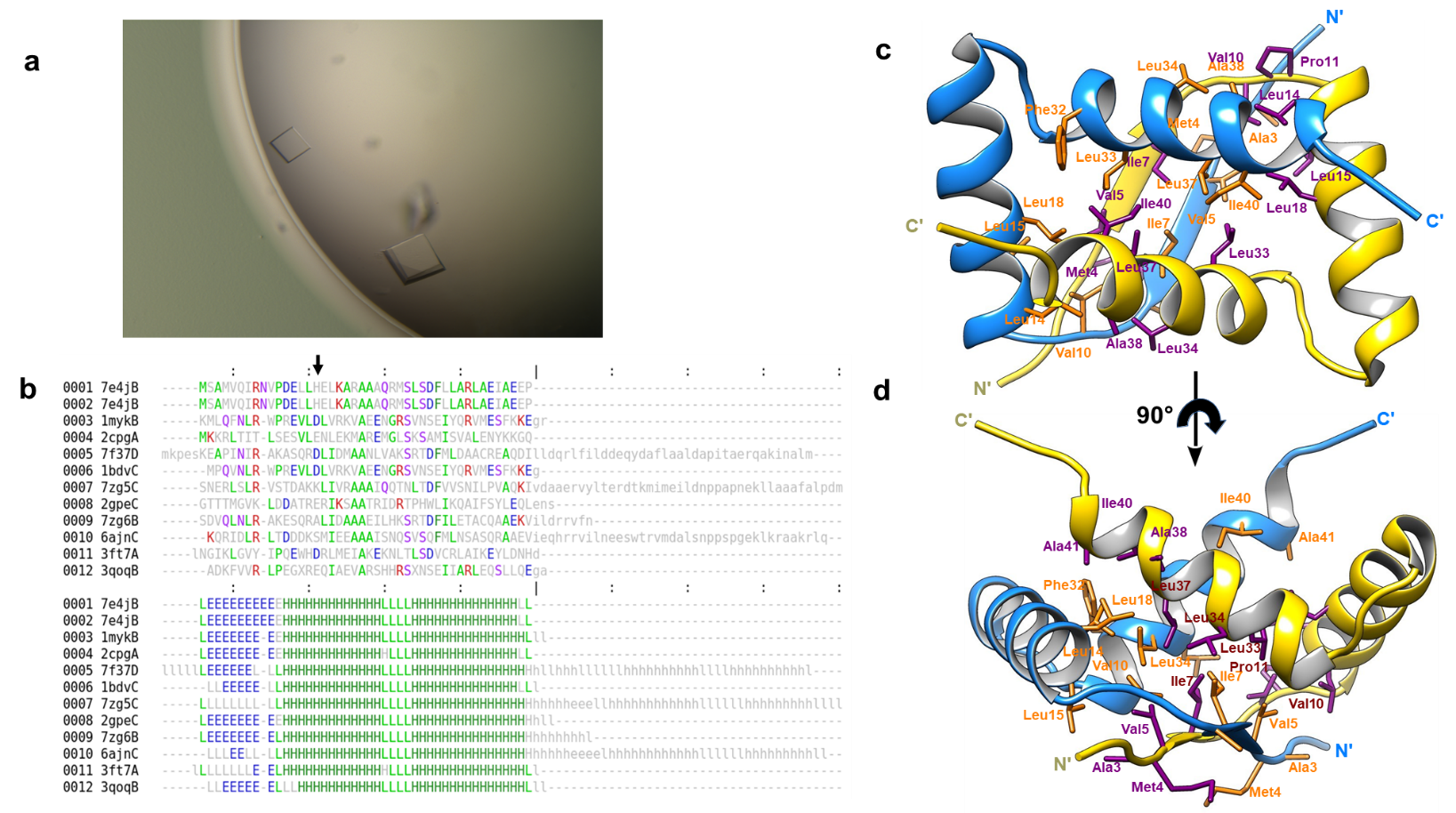

**Fig.S2 : Detailed structures of the VapB12 homodimeric crystal structure interface.** (a). VapB12 protein crystal grown in 0.2 M ammonium chloride-NaOH pH 6.3, 20% (w/v) PEG 3350. (b). Structure-based sequence alignment of VapB12 with its top ten structural homologs identified by a Dali search. Location of His16 position in VapB12 (PDB ID: 7E4J) is indicated by a black arrow at the top. The first part (top) shows the amino acid sequences of the structural neighbors, while the second part (bottom) shows the secondary structure assignments by DSSP (H/h: helix, E/e: strand, and L/l: coil). The most frequent amino acid type is colored in each chain. (c) Ribbon representation of the structure of VapB12 homodimer. Residues involved in the formation of the hydrophobic interface from both subunits are represented as sticks labeled with residue names and numbers rendered in orange and violet colors. (d) is the view of 90° rotation of (c) along the x-axis.

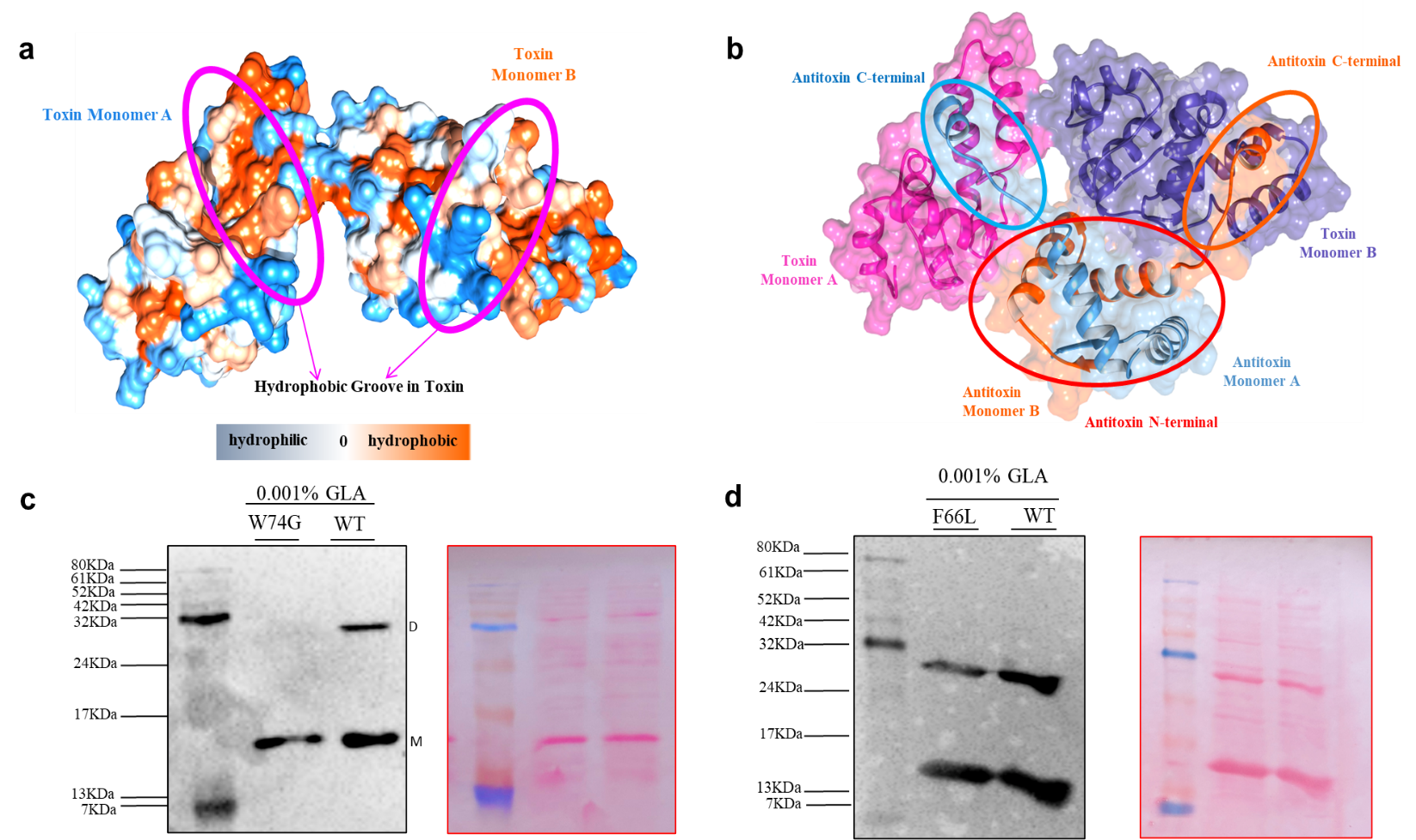

**Fig.S3 : Structural basis of the VapBC12 Toxin-Antitoxin complex formation**. (a) A hydrophobicity map of the VapC12 toxin dimer's molecular surface, ranging from hydrophilic (blue) to hydrophobic (orange). This reveals two deep, hydrophobic grooves on the dimer that form the primary interface for antitoxin recognition. (b) The structural model of the VapC12-VapB12 heterotetrameric complex. The antitoxin C-terminal helices (shown in light blue and orange, highlighted by the red oval) insert into the corresponding hydrophobic grooves of the toxin monomers (magenta and purple). This interaction is critical for neutralizing the toxin's activity. The globular N-terminal domains of the antitoxin monomers bind to distal sites on the toxin. (c-d): complete image of western blots in Fig 3e & g and their ponceau image at the respective right side. Band sizes of ladder (pink plus prestained protein ladder) are mentioned on the right side of the blot image. M, monomer = 14KDa. D, dimer = 28KDa.

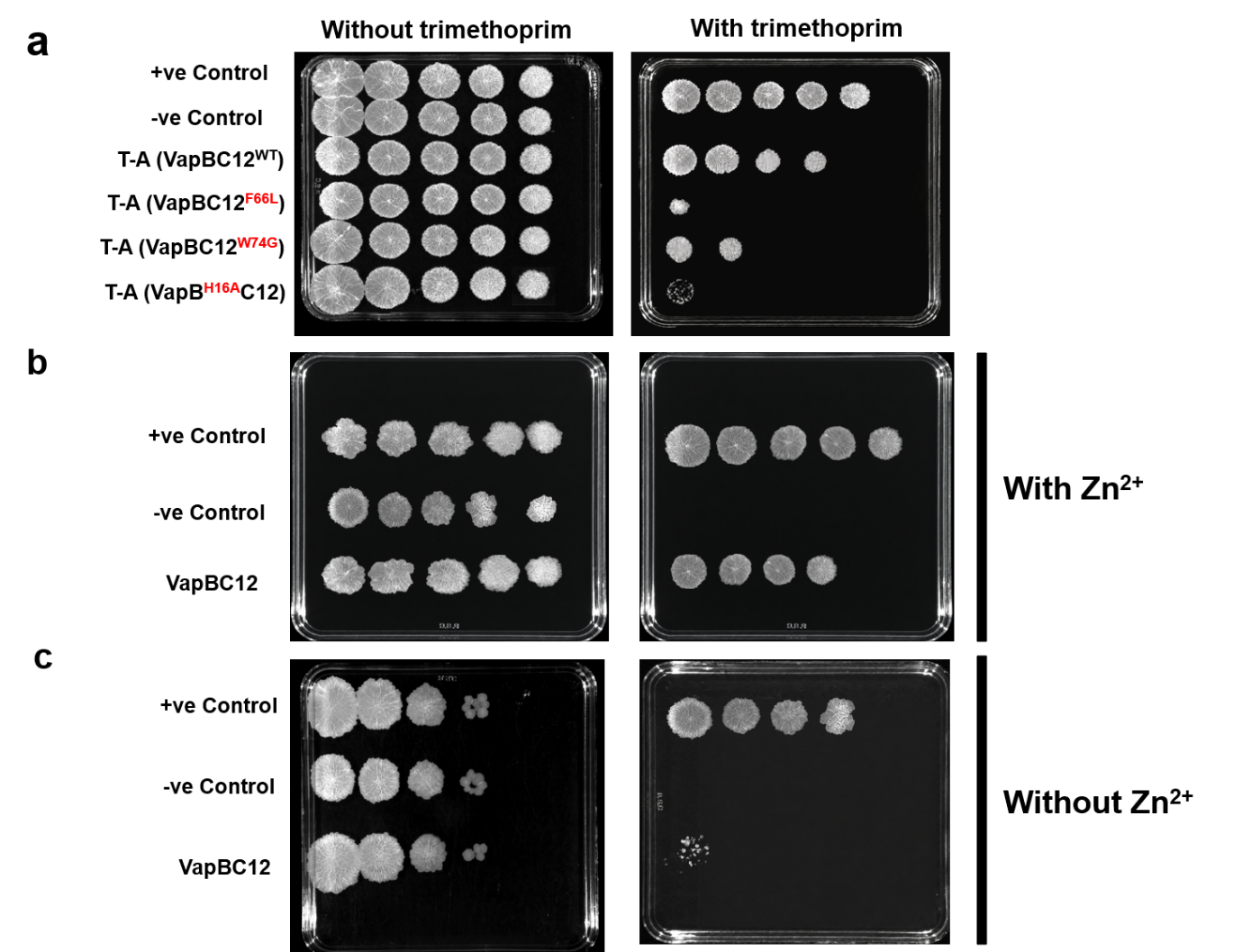

**Fig.S4 : Interaction analysis of VapBC12 proteins using MPFC assay. (a)** Complete image of the agar plates showing ten-fold serial dilution spot titers of the indicated strains in the presence or absence of trimethoprim. The plate images were used to generate the spot-titer plots shown in Fig. 3g and Fig. 4a,b. **(b-c)** Agar plates supplemented with trimethoprim, with **(b)** or without zinc sulfate **(c),** showing the interaction between VapBC12-containing strains compared with the positive and negative controls. The images were used to generate Fig 4b,c.

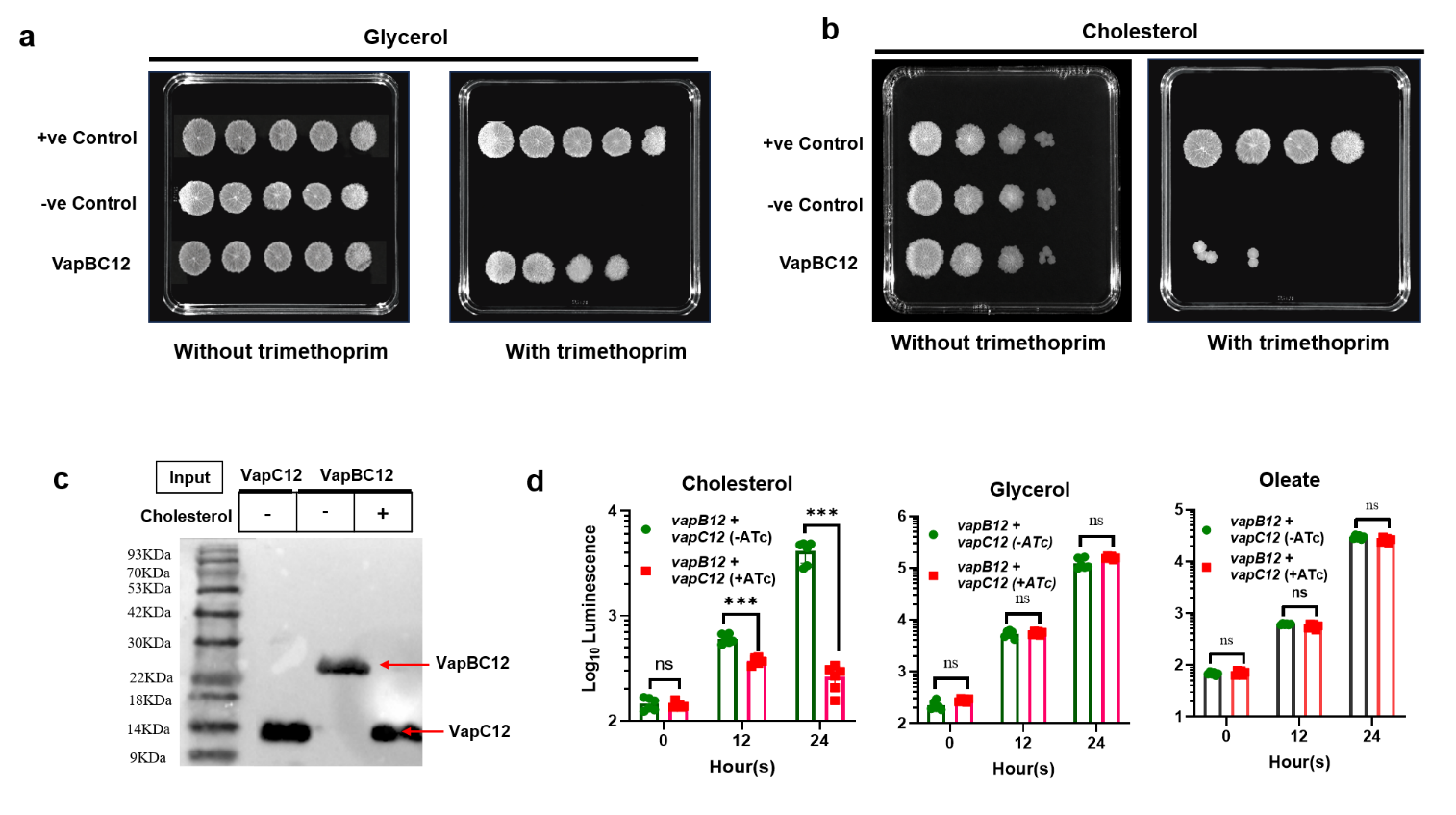

**Fig.S5 : Interaction analysis of VapBC12 proteins in presence of Cholesterol. (a-b)** Complete image of the agar plates showing ten-fold serial dilution spot titers of the indicated strains with and without trimethoprim supplemented with Glycerol **(a)** and Cholesterol **(b)**. The plate images were used to generate the spot-titer plots shown in Fig. 5c. **(c)** Complete image of western blot in Fig 5d. Band sizes of ladder (BlueAQUA Prestained Protein Ladder) are mentioned on the right side of the blot image. **(d)** Time dependent growth analysis for figure presented in Fig 5e. Luminescence was measured at indicated time points.

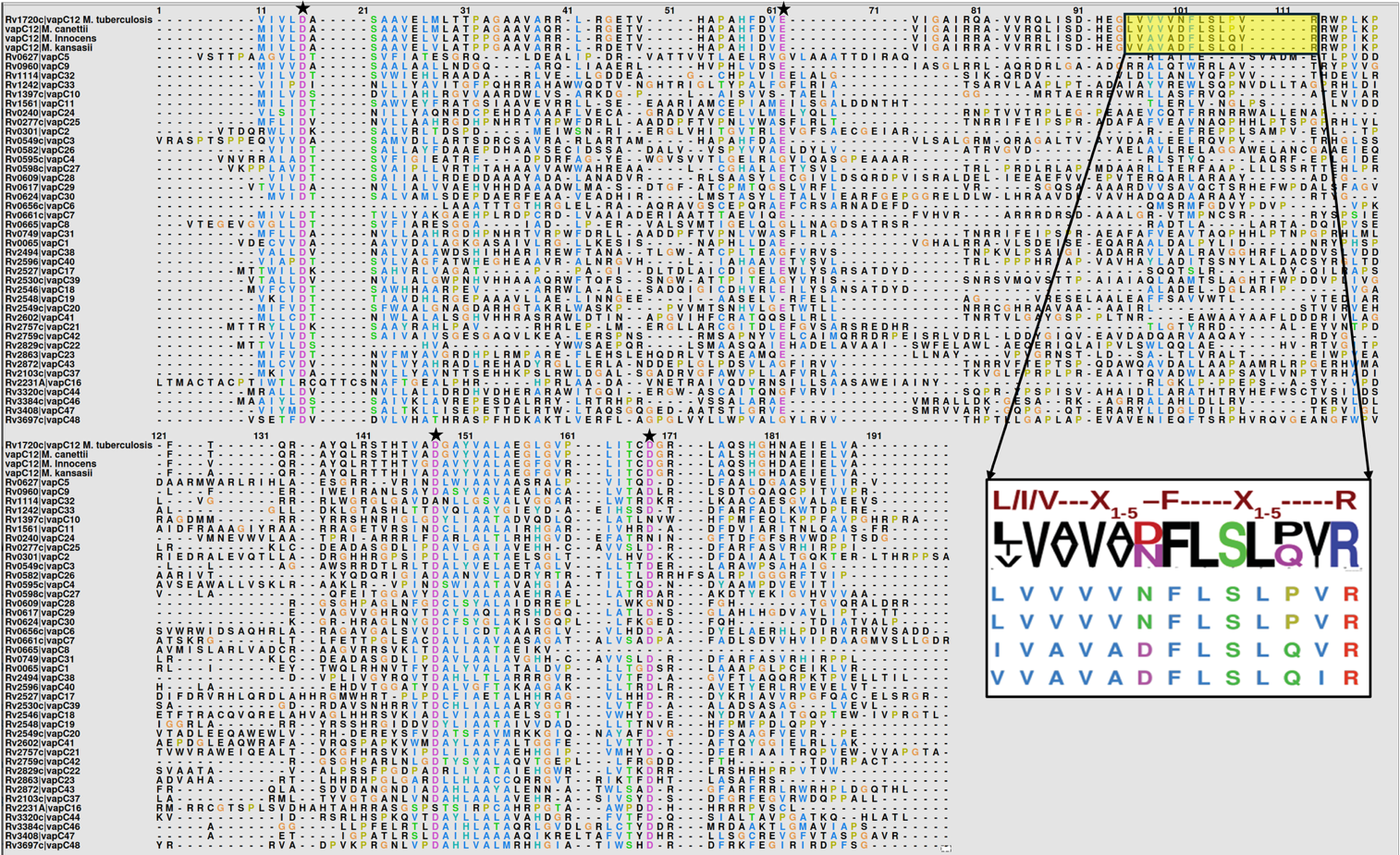

**Fig.S6 : CRAC motif is unique to VapC12 toxin of *M. tuberculosis:*** Multiple Sequence Alignment (MSA) of VapC12 toxin with other VapC class toxins from Mtb and VapC12 from other mycobacterial species (e.g., *M. canettii, M.innocens, M. kansasii*). The alignment was generated using Clustal Omega and visualized using Chimera. The cholesterol-recognition amino acid consensus (CRAC) motif ([L/I/V]-X1–5-[F]-X1–5-[K/R]), a conserved cholesterol-binding domain, is highlighted in a box. The conserved active sites and magnesium-binding residues are indicated with an asterisk (*) at the top. The numbers at the top represent the residue positions. Sequence conservation is denoted by shading intensity (see key). The enlarged motif inset (bottom) was generated using Sequence Logo to emphasize the residue frequency and conservation.

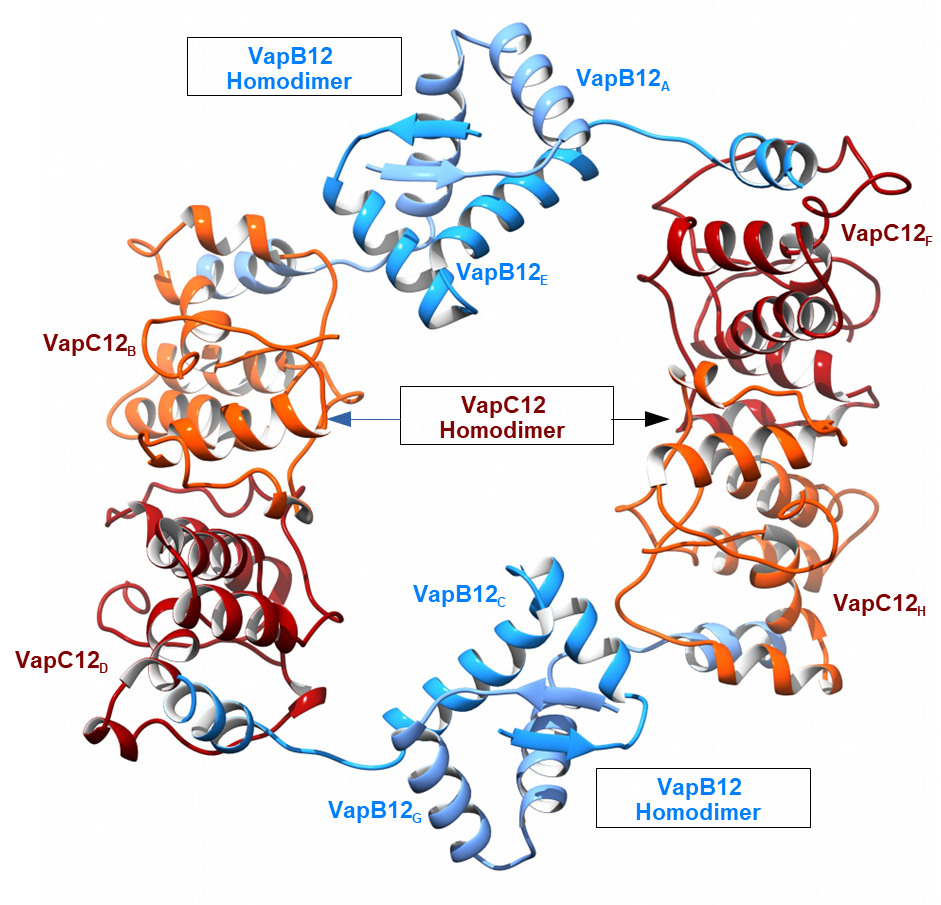

**Fig.S7 : Overall structure of the homology docked model of VapBC12 and its subunits.** Ribbon representation of the VapBC12 hetero-octamer model obtained by homology docking. Chains G and E of VapB12 are shown in blue and chains C and A are shown in sky-blue. Chains B and H and chains D and F of VapC12 are shown in orange and red, respectively.

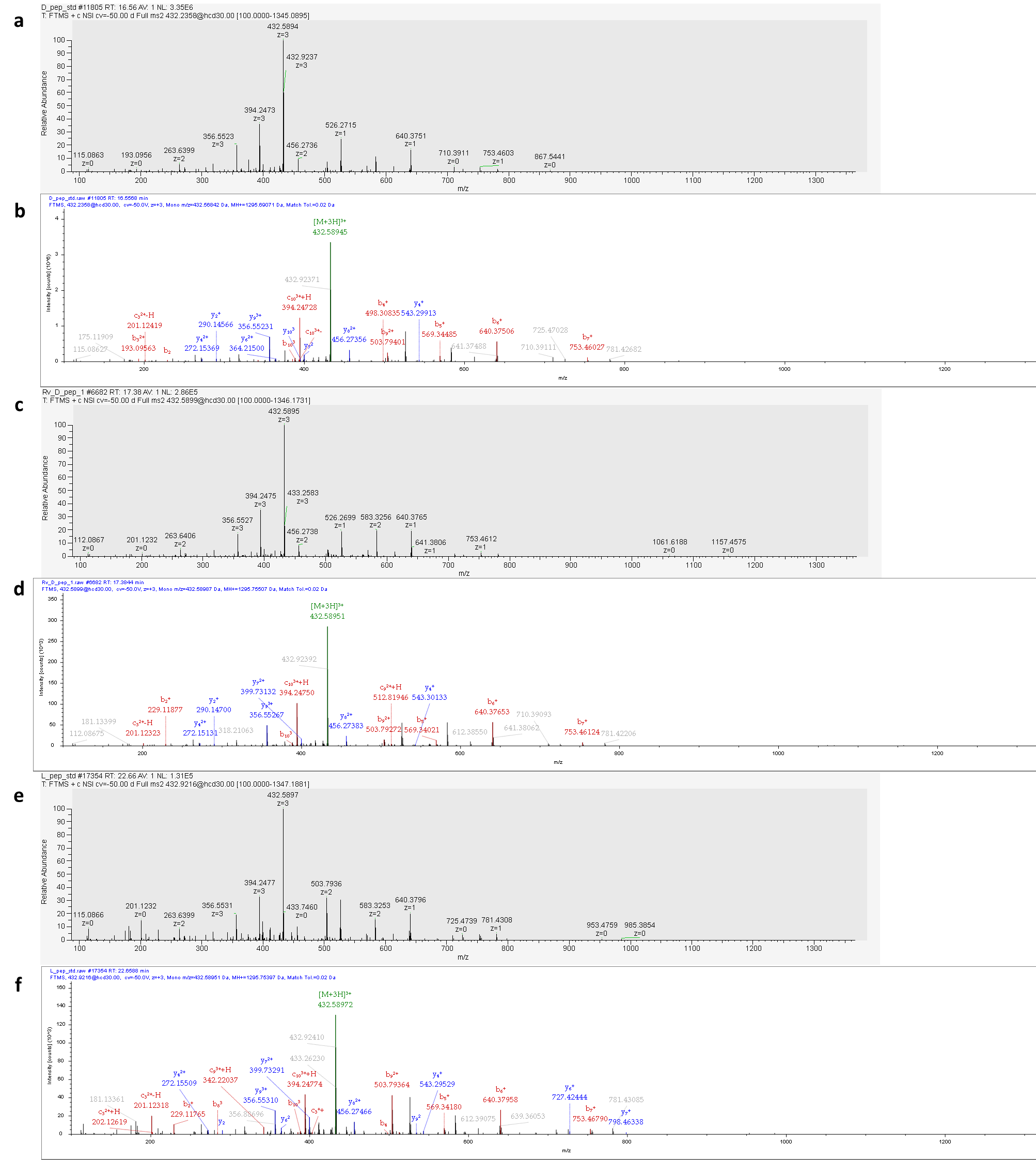

**Fig.S8 :** **Intracellular stability and recovery of D- and L-enantiomers of the P2 peptide in *M. tuberculosis* by LC–MS/MS.** Representative MS/MS spectra of **(a)** raw and **(b)** annotated (by Proteome Discoverer) purified D-P2 peptide standard, **(c)** raw and **(d)** annotated intact D-P2 peptide recovered from whole-cell lysates of *M. tuberculosis* treated with 10 µM D-P2 for 4 days (starting OD_600_ 0.01), and **(e)** raw and **(f)** annotated purified L-P2 peptide standard. Whole-cell lysates from *M. tuberculosis* cultures treated in parallel with 10 µM L-P2 for 4 days were analyzed under identical LC–MS/MS conditions; no precursor or fragment ions corresponding to unmodified L-P2 were detected above background, and therefore no fragmentation spectrum was acquired.

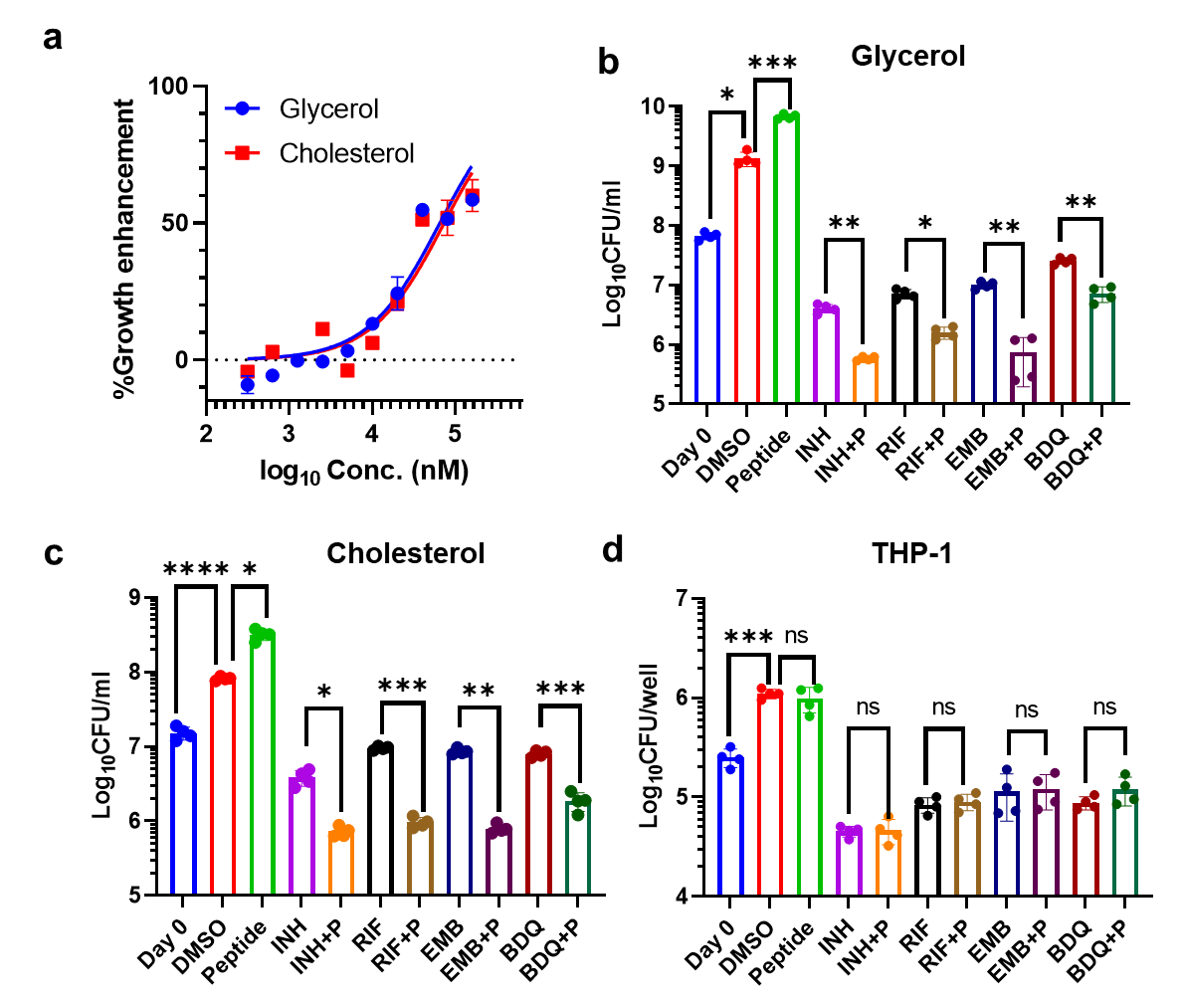

**Fig.S9 : Activity of L-form P2 peptide as an adjunct to antibiotic treatment in *M. tuberculosis*. (a)** Dose–response analysis of growth enhancement in glycerol- and cholesterol-containing media in the presence of L-form P2 peptide normalised with DMSO control. EC₅₀ and EC₉₉ values were calculated by non-linear regression using GraphPad Prism. Data represent mean ± SEM from two experiments with technical replicates. **(b-c)** *In vitro* antibiotic killing of H37Rv with 10× MIC isoniazid (INH), rifampicin (RIF), ethambutol (EMB), or bedaquiline (BDQ), alone or combined with 38 µM L-form P2, in glycerol (a) or cholesterol (b) media. **(c)** Intracellular survival in THP-1–derived macrophages measured by CFU enumeration following infection (MOI 1:1) and treatment with 1× MIC antibiotics ± 40 µM L-form P2. Data are mean ± SD of two biological replicates. Two-way ANOVA was used for data analysis; ns, not significant; **p < 0.01; ***p < 0.001; ****p < 0.0001

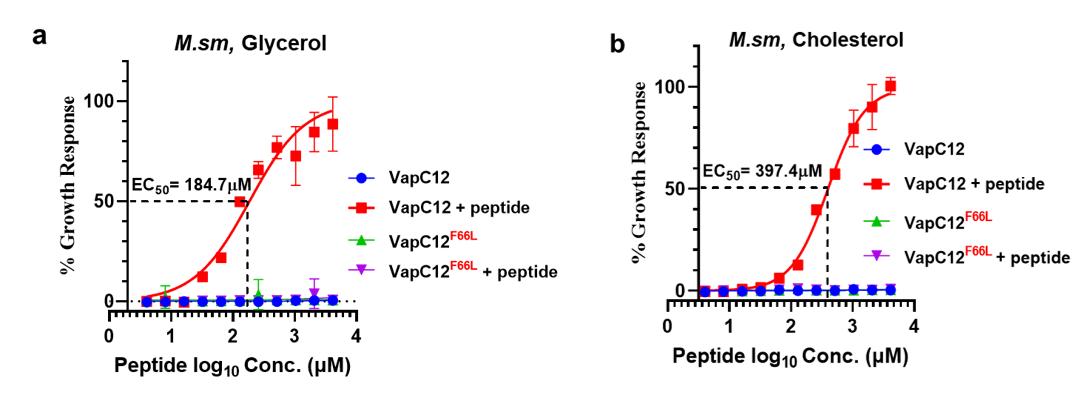

**Fig.S10: *In vitro* activity of P2 peptide.** **(a-b)** P2 peptide dose-response curve (DRC) against WT-VapC12 and mutated VapC12^F66L^ toxin activity: strain expressing *vapC12* and *vapC12^F66L^* gene was incubated at density of 10^5^ cells/ml in a 96-well white plate with and without P2 peptide in glycerol (0.1%) and cholesterol (150µM) Media. The peptide was two-fold serially diluted to obtain minimum effective concentration (MEC). The black dashed line marks the EC50 (184.7 µM) in glycerol (a)- media and (397.4 µM) cholesterol (b) media. EC₅₀ values were calculated by non-linear regression using GraphPad Prism.

**Supplementary Table 1: List of vectors used in the study**

| **S. No.** | **Plasmid** | **Feature** | **Reference** |
| --- | --- | --- | --- |
| 1. | pST-K | Kan, episomal, mycobacterial expression vector | Kind gift from Dr. Vinay  K. Nandicoori, CSIR-CCMB, India |
| 2. | pST-KT | Kan, *tet-*inducible, mycobacterial expression vector | Kind gift from Dr. Vinay  K. Nandicoori, CSIR-CCMB, India |
| 3. | pST-H | Hyg, pST-K backbone with hyg-cassette | Kind gift from Dr. Vinay  K. Nandicoori, CSIR-CCMB, India |
| 4. | pET28a | Kan, IPTG-inducible, *E.coli* with expression vector | This lab |
| 5. | pMD101 | Kan, Mycobacterial M-PFC system | Kind gift from Dr. Ashwani Kumar, CSIR-IMTECH, India |
| 6. | pMD102 | Hyg, Mycobacterial M-PFC system | Kind gift from Dr. Ashwani Kumar, CSIR-IMTECH, India |
| 7. | pBP10 | Kan, episomal, replication clock plasmid | Kind gift from Prof. David Sherman |

**Supplementary Table 2: Strains used in M-PFC (PPI) study**

| **S.No.** | **Strain** | **Bait** | **Prey** | **Selection Marker** |
| --- | --- | --- | --- | --- |
| 1. | pMD101+pMD102 | pMD101 | pMD102 | Kan+Hyg |
| 2. | pMD101:CFP10  +  pMD102-ESAT-6 | pMD101: CFP10 | pMD102:ESAT6 | Kan+Hyg |
| 3. | pMD101-VapC12  +  pMD102:VapB12 | pMD101VapC12 | pMD102:VapB12 | Kan + Hyg |
| 4. | pMD101:VapC12 (F66L)  +  pMD102:VapB12 | pMD101:VapC12  (F66L) | pMD102:VapB12 | Kan + Hyg |
| 5. | pMD101:VapC12 (W74G)  +  pMD102:VapB12 | pMD101:VapC  (W74G) | pMD102:VapB | Kan + Hyg |
| 6. | pMD101:VapC12  +  pMD102:VapB12 (H16A) | pMD101:VapC | pMD102:VapB (H16A) | Kan + Hyg |

**Supplementary Table 3: List of bacterial strains used in the study**

| **S.No.** | **Strain (Host)** | **Feature/Purpose** | **Source/Reference** |
| --- | --- | --- | --- |
| 1. | *E. coli* XL1Blue | Plasmid storage and amplification | This lab |
| 2. | *E. coli* BL21 codon+ | Heterologous Protein expression | This lab |
| 3. | *M. smegmatis*  mc^2^155 | Wild type *M. smegmatis* / Growth Kinetics | Kind gift from Sarah Fortune, Harvard T.H. Chan School of Public Health |
| 4. | *M. tuberculosis*  H37Rv | Wild type *M. tuberculosis /* Growth Kinetics | Kind gift from Christopher M.Sassetti, UMass |
| 5. | *ΔVapC12* | *Rv1720*, *VapC12* deletion mutant, unmarked | This lab |
| 6. | pSTK:VapC12 (*M. smegmatis*) | Growth Kinetics | This study |
| 7. | pSTK (*M. smegmatis*) | Growth Kinetics | This study |
| 8. | pSTKT:VapC12 (*M. smegmatis*) | Growth Kinetics | This study |
| 9. | pSTK:VapC12 (W74G) (*M. smegmatis*) | Growth Kinetics | This study |
| 10. | pSTK:VapC12 (F66L)  (*M. smegmatis*) | Growth Kinetics | This study |
| 11. | pSTKT:VapC12  + pSTH:VapB12 (*M. smegmatis*) | Growth Kinetics | This study |
| 12. | pSTK:VapBC12 (*M. smegmatis*) | Growth Kinetics | This study |
| 13. | pSTK:VapBC12 (F66L) (*M. smegmatis*) | Growth Kinetics | This study |
| 14. | pSTK:VapBC12(W74G)  (*M. smegmatis*) | Growth Kinetics | This study |
| 15. | pSTK:VapB(H16A) C12 (*M. smegmatis*) | Growth Kinetics | This study |
| 16. | pET22b: VapBC12 (E.coli, BL21 codon+) | Protein expression | This study |
| 17. | pET28a:VapC12 (E.coli, BL21 codon+) | Protein expression | This study |
| 18. | pET28a:VapC12 (W74G) (E.coli, BL21 codon+) | Protein expression | This study |
| 19. | pET28a:VapC12 (F66L) (E.coli, BL21 codon+) | Protein expression | This study |
| 20. | pET22b (E.coli, BL21 codon+) | Protein expression | This study |
| 21. | pMD101+pMD102 (*M. smegmatis*) | Protein-Protein Interaction | Kind gift from Dr. Ashwini Kumar, IMTech, Chandigarh |
| 22. | pMD101: CFP10 + pMD102: ESAT-6 (*M. smegmatis*) | Protein-Protein Interaction | Kind gift from Dr. Ashwini Kumar, IMTech, Chandigarh |
| 23. | pMD101:VapC12 + pMD102: VapB12 (*M. smegmatis*) | Protein-Protein Interaction | This study |
| 24. | pMD101:VapC12 (F66L) + pMD102: VapB12 (*M. smegmatis*) | Protein-Protein Interaction | This study |
| 25. | pMD101:VapC12 (W74G) + pMD102: VapB12 (*M. smegmatis*) | Protein-Protein Interaction | This study |
| 26. | pMD101:VapC12 + pMD102: VapB12 (H16A) (*M. smegmatis*) | Protein-Protein Interaction | This study |

**Supplementary Table 4: List of Primers used in this study**

| **S.No.** | **Primers** | **Sequence** | **RE used** | **Purpose** |
| --- | --- | --- | --- | --- |
| 1. | VapC12 FP’ | ATTGGATCCGTGATCGTGTTGGACGCCTC | BamH1 | Cloning for Growth Kinetics, Protein expression, PPI |
| 2. | VapC12 RP’ | ATTGAATTCTCAGGCGACAAGCTCGATCTC | EcoR1 | Cloning for Growth Kinetics, Protein expression, PPI |
| 3. | VapBC12 FP’ | ATACATATGATGTCCGCCATGGTTAGATCC | Nde1 | Cloning for Growth Kinetics, Protein expression |
| 4. | VapBC12 RP’ | AATAAGCTTGGCGACAAGCTCGACTCC | HindIII | Cloning for Growth Kinetics, Protein expression |
| 5. | VapC12(F66L) FP’ | TCAACTTGCTTAGCCTGCCCG |  | SDM generation for Growth Kinetics, Protein expression, PPI |
| 6. | VapC12 (F66L) RP’ | GCTAAGCAAGTTGACAACCAGGAG |  | SDM generation for Growth Kinetics, Protein expression, PPI |
| 7. | VapC12 (W74G) FP’ | CGTTCGGCGTGAGCCGT |  | SDM generation for Growth Kinetics, Protein expression, PPI |
| 8. | VapC12 (W74G) RP’ | GCTTTAACGGCTCACGCC |  | SDM generation for Growth Kinetics, Protein expression, PPI |
| 9. | VapB12 FP’ | ATTGGATCCATGTCCGCCATGGTTCAGATCC | BamH1 | Cloning for Growth Kinetics, Protein expression, PPI |
| 10. | VapB12 RP’ | ATTGAATTCTCACTCAGATCGAGCCTCGTC | EcoR1 | Cloning for Growth Kinetics, Protein expression, PPI |
| 11. | VapB12 (H16A) FP’ | GAGCTTCTCGCCGAGCTG |  | SDM generation for Growth Kinetics, Protein expression, PPI |
| 12. | VapB12 (H16A) RP’ | GCCTTCAGCTCGGCGAGA |  | SDM generation for Growth Kinetics, Protein expression, PPI |
